# Synaptotagmins Maintain Diacylglycerol Homeostasis at Endoplasmic Reticulum-Plasma Membrane Contact Sites during Abiotic Stress

**DOI:** 10.1101/2020.07.28.222919

**Authors:** Noemi Ruiz-Lopez, Jessica Pérez-Sancho, Alicia Esteban del Valle, Richard P. Haslam, Steffen Vanneste, Rafael Catalá, Carlos Perea-Resa, Daniël Van Damme, Selene García-Hernández, Armando Albert, José Vallarino, Jinxing Lin, Jir□í Friml, Alberto P. Macho, Julio Salinas, Abel Rosado, Johnathan A. Napier, Vitor Amorim-Silva, Miguel A. Botella

## Abstract

Endoplasmic Reticulum-Plasma Membrane contact sites (ER-PM CS) play fundamental roles in all eukaryotic cells. Arabidopsis mutants lacking the ER-PM protein tether synaptotagmin1 (SYT1) exhibit decreased plasma membrane (PM) integrity under multiple abiotic stresses such as freezing, high salt, osmotic stress and mechanical damage. Here, we show that, together with *SYT1*, the stress-induced *SYT3* is an ER-PM tether that also functions in maintaining PM integrity. The ER-PM CS localization of SYT1 and SYT3 is dependent on PM phosphatidylinositol-4-phosphate and is regulated by abiotic stress. Lipidomic analysis revealed that cold stress increased the accumulation of diacylglycerol at the PM in a *syt1/3* double mutant relative to WT while the levels of most glycerolipid species remain unchanged. Additionally, SYT1-GFP preferentially binds diacylglycerol *in vivo* with little affinity for polar glycerolipids. Our work uncovers a crucial SYT-dependent mechanism of stress adaptation counteracting the detrimental accumulation of diacylglycerol at the PM produced during episodes of abiotic stress.

## INTRODUCTION

As sessile organisms, plants are continuously exposed to changes in environmental conditions (Botella et al., 2007; Zhu, 2016). In addition to form a selective permeable barrier, the plasma membrane (PM) senses environmental cues and transforms them into highly regulated signaling outputs (Hou et al., 2016). Among these outputs, there are several lipids such as phosphatidic acid (PA) and phosphatidylinositol phosphates (also known as phosphoinositides, PIPs), which are synthetized at the PM and play important regulatory roles in cellular plasticity (Colin and Jaillais, 2020; Hou et al., 2016; Munnik and Nielsen, 2011; Munnik and Testerink, 2009; Testerink and Munnik, 2005). In particular, PA is rapidly generated at the PM in response to abiotic stresses such as dehydration (Hong et al., 2008), salinity (Munnik et al., 2000; L. Yu et al., 2010), temperature stress (Arisz et al., 2013; Ruelland, 2002) and treatment with the stress-related hormone abscisic acid (ABA) (Katagiri et al., 2005; Li et al., 2019; Uraji et al., 2012). The production of PA at the PM can be triggered by the hydrolysis of structural phospholipids such as phosphatidylcholine (PC) and phosphatidylethanolamine (PE), either directly by phospholipases D (PLD) or through the consecutive actions of non-specific phospholipases C (NPC), that produce diacylglycerol (DAG), which is transformed into PA by diacylglycerol kinases (DGKs) (Pokotylo et al., 2018). In addition, hydrolysis of the PIPs, e.g. phosphatidylinositol-4-phosphate (PI4P) and phosphatidylinositol-4,5-biphosphate (PI4,5P_2_), by phospholipase C (PLC) also produces DAG, that can be transformed into PA by DGKs (Arisz et al., 2009; Testerink and Munnik, 2011) (Arisz et al., 2013). Because DAG accumulation causes the disruption of the lamellar phase of lipid membranes, its concentration must be tightly controlled (Campomanes et al., 2019).

The PM forms extensive contacts with the endoplasmic reticulum (ER) at Endoplasmic Reticulum-Plasma Membrane Contact Sites (ER-PM CS) which can be defined as tight and not fused membrane junctions tethered by specific protein complexes (Bayer et al., 2017; Scorrano et al., 2019; Wu et al., 2018). These ER-PM CS are essential for communication between the ER and PM in mammalian, fungal and plant cells, enabling lipid transport (Saheki et al., 2016), regulating calcium influx (Saheki and De Camilli, 2016), and maintaining the cortical ER morphology (Siao et al., 2016). The plant Synaptotagmins (SYTs) and their ortholog counterparts, the mammalian Extended Synaptotagmins (E-Syts) and the yeast Tricalbins (Tcbs) are ER-PM CS tethers that share a common modular structure, comprising a N-terminal transmembrane (TM) domain that anchors them to the ER, a synaptotagmin-like mitochondrial-lipid binding (SMP) domain, and a variable number of the Ca^2+^- and phospholipid-binding C2 domains (Giordano et al., 2013; Manford et al., 2012; Perez Sancho et al., 2015). SMP domains are members of the <u>tu</u>bular <u>li</u>pid-binding <u>p</u>rotein (TULIP) superfamily, with a common folding structure that can harbor lipids in a hydrophobic cavity (Kopec et al., 2010). The modular structure of these proteins implies two likely inter-related functions, namely the establishment of Ca^2+^ regulated ER-PM tethering by their C2 domains (Giordano et al., 2013; Manford et al., 2012; Perez Sancho et al., 2015; Saheki et al., 2016), and the SMP dependent transport of lipids between the PM and the ER (Saheki et al., 2016; Schauder et al., 2014).

The yeast Tcbs and the mammalian E-Syts contain a variable number of C2 domains, which can bind membrane phospholipids either dependent or independently of Ca^2+^. In yeast, the C2C domains of Tcb1 and Tcb3 bind anionic phospholipids in a Ca^2+^-dependent manner, while all the C2 domains in Tcb2 are insensitive to Ca^2+^ (Schulz and Creutz, 2004). In mammals, E-Syt2 and E-Syt3 are localized at ER-PM CS due to the constitutive interaction of their C2 domains with PM PI(4,5)P_2_ meanwhile E-Syt1 is localized throughout the ER at resting state (Fernández-Busnadiego et al., 2015; Giordano et al., 2013; Idevall-Hagren et al., 2015). However, elevation of cytosolic Ca^2+^ promotes its accumulation at the ER–PM CS and reduces the distance between ER and PM (Bian et al., 2018; Fernández-Busnadiego et al., 2015; Giordano et al., 2013; Idevall-Hagren et al., 2015). Interestingly, loss of all three Tcbs does not result in a substantial reduction of ER-PM CS in yeast (Manford et al., 2012), and suppression of all three E-Syts in mice does not affect their normal development, viability or fertility nor their ER morphology (Sclip et al., 2016; Tremblay and Moss, 2016), indicating the presence of additional tethers.

A recent study has shown that Tcbs are necessary for the generation of peaks of extreme curvature at cortical ER (cER) membrane and the maintenance of PM integrity in yeast. Although the study proposed that these processes were facilitated by the transport of lipids from the highly curved, less packaged, cER to the PM (Collado et al., 2019), the nature of the lipids transported remains elusive. Important insights about the function of E-Syts was provided by the structural analyses of the SMP domain of E-Syt2 (Schauder et al., 2014). The SMP-domain of E-Syt2 consists of six beta strands and two alpha helices arranged in a barrel that homodimerizes to form a cylinder whose interior is lined almost exclusively by hydrophobic residues. Importantly, the crystal structure of the SMP dimer revealed the presence of glycerophospholipids in the hydrophobic channel without preference for any particular headgroup (Schauder et al., 2014). Further studies indicated that E-Syt1 also transfers glycerolipids *in vitro* without any preference for the head group and that an E-Syt1 mutant lacking the SMP domain lacks the lipid transfer ability (Saheki et al., 2016; H. Yu et al., 2016). Despite the difficulty of identifying the specific lipids preferentially transported by the SMP domain of E-Syts, analysis of cells lacking all three E-Syts supports the hypothesis that the E-Syts function *in vivo* is to clear DAG formed at the PM after receptor-triggered activation of phospholipase C (PLC) (Bian et al., 2018; Saheki et al., 2016). This proposed role of E-Syts in DAG homeostasis has been further supported by the finding that E-Syt1 regulates insulin secretion in pancreatic islets by clearing DAG from the PM (Xie et al., 2019).

The Arabidopsis genome encodes five SYTs (SYT1-SYT), all of them containing a TM region, an SMP domain and two C2 domains (Perez Sancho et al., 2016). The C2 domains of SYT1 are targeted to the PM through their binding to negatively charged PIP, resulting in a constitutive localization of SYT1 at ER-PM CS (Perez Sancho et al., 2015). Originally, Arabidopsis *SYT1* was identified using a forward genetic screen based on salt hypersensitivity (Schapire et al., 2008) and as a cold-induced protein required for Ca^2+^-dependent freezing tolerance (Yamazaki et al., 2008). Interestingly, salt stress causes an increase of SYT1 localization at ER-PM CS as well as an expansion of these ER-PM CS (Lee et al., 2019). In the last few years, additional roles for SYT1 have been uncovered based on the analysis of loss-of-function *syt1* mutants, including the stability of cER network (Ishikawa et al., 2018; Siao et al., 2016), wounding responses (Perez Sancho et al., 2015) and the resistance to several biotic factors such as viruses (Levy et al., 2015; Lewis and Lazarowitz, 2010; Uchiyama et al., 2014) and fungi (Kim et al., 2016). Recently, it has been shown that SYT5, a major interactor of SYT1, also localizes at ER-PM CS (Ishikawa et al., 2020; Lee et al., 2020) and displays similar changes in localization in response to salt stress as SYT1 (Lee et al., 2020).

Despite the defective responses reported for *syt1* under multiple biotic and abiotic stresses a mechanistic understanding is still lacking. It has been long established that abiotic stresses produce lipid changes at the PM, therefore we hypothesized that SYT1 may function in the homeostasis of PM lipid composition by transporting lipids at ER-PM CS and as a consequence, in the tolerance for multiple and apparently unrelated stresses. Here, we reveal that SYT1 and SYT3 are ER-PM tethers that play an important role in PM lipid homeostasis by transporting DAG that is produced at the PM as a consequence of different abiotic stresses. Additionally, monitored dynamics of SYT1 and SYT3 in responses to stress are consistent with their role in lipid transport in order to maintain PM integrity under adverse environmental conditions.

## RESULTS

### SYT1 and SYT3 are important for PM stability under abiotic stress

Earlier work has established that the *SYT1* gene is an important determinant of cell viability under various abiotic stresses (Perez Sancho et al., 2015; Schapire et al., 2008; Yamazaki et al., 2008), therefore we investigated whether other *SYTs* might also have a role in stress resistance. The Arabidopsis genome encodes five SYT homologues that can be divided into two clades (Figure S1A). *In silico* analysis of *SYT* genes in vegetative tissues using available RNA sequencing data, e.g. eFP Browser (Figure 1A) and TRAVA (Figure S1B) showed that *SYT1, SYT3*, and *SYT5* are ubiquitously expressed with *SYT1* being about 6 and 10 times more expressed than *SYT5* and *SYT3*, respectively. *SYT4* and *SYT2*, however, have low expression in vegetative tissues. Expression analysis of abiotic stress responses of *SYT* genes using available transcriptomic databases indicated a stress regulated expression for *SYT1* and *SYT3* but not *SYT5* (Figure S1C, S1D). Consistently, when we experimentally analyzed the cold response of all *SYT* genes at the transcriptional level using RT-qPCR, *SYT1* and *SYT3* but not *SYT2, SYT4* or *SYT5* were induced after 24 hr of cold stress (Figure 1B). Thus, we focused on SYT1 and SYT3 to further characterize their involvement in cell viability under abiotic stress conditions.

**Figure 1.**
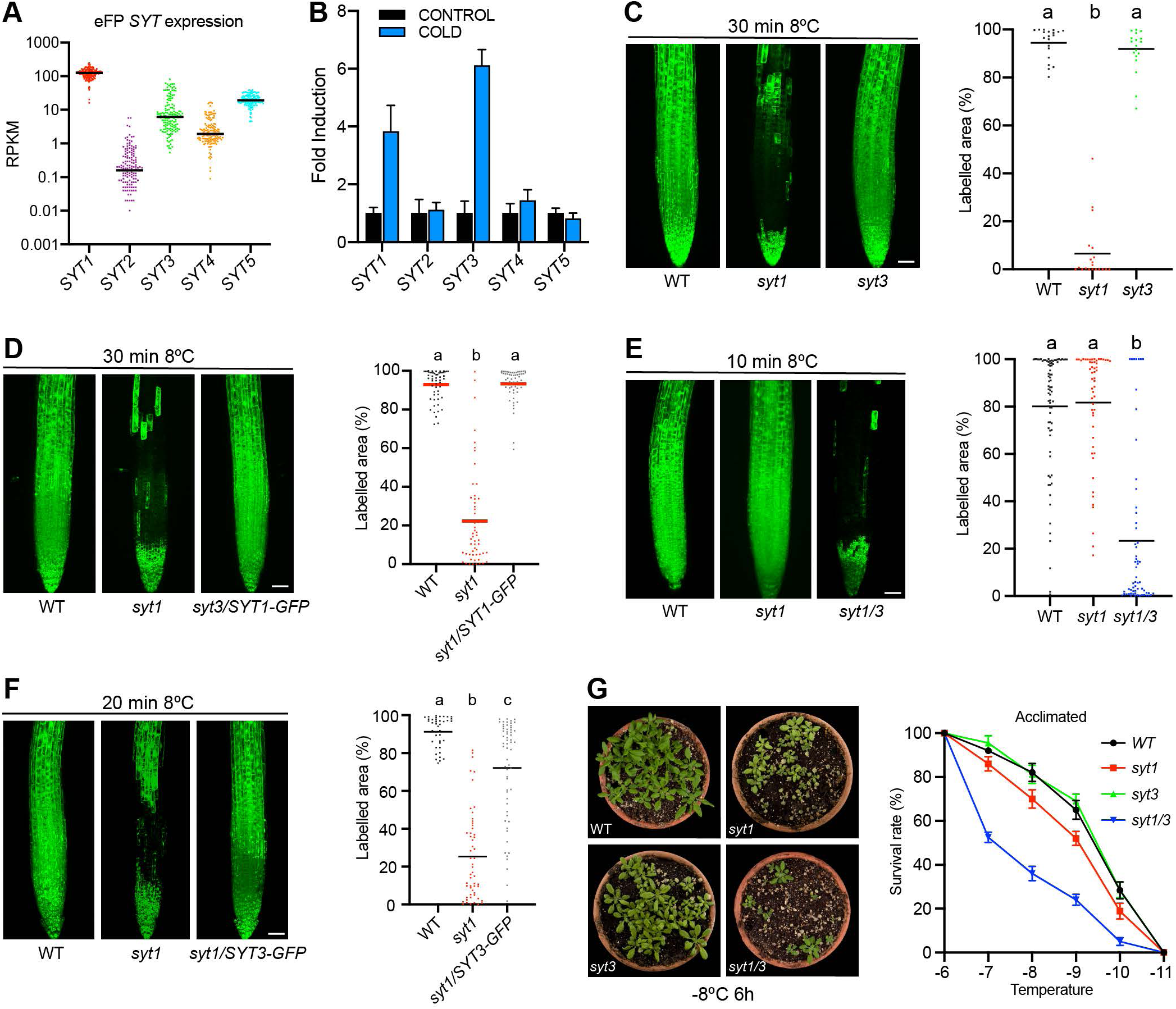
SYT1 and SYT3 are important for PM stability under abiotic stress. RNA-seq data of *SYT1, SYT2, SYT3, SYT4*, and *SYT5* obtained from vegetative tissues at different developmental stages from eFP-seq Browser (https://bar.utoronto.ca/eFP-Seq_Browser/). Each dot represents a value of RPKM and the bar represent the median. *SYT1* and *SYT3* transcripts are induced by cold. Arabidopsis WT seedlings were grown under long-day photoperiod and 23°C for 7 days and then transferred to 4°C for 24 h or kept under control conditions. The relative expression level of the *SYT* genes was measured by RT-qPRC and normalized to the expression of *ACTIN2*. Error bars indicate SD. (C to F) (C to F) Confocal images and cell viability quantification after cold treatment in 6-day-old Arabidopsis roots roots. Seedlings grown in half-strength MS agar solidified medium under long-day photoperiod and 23°C. Cell viability was determined by fluorescein diacetate (FDA) staining and quantified as the percentage of root area with fluorescence above an automatic threshold stablished by the “Moments” algorithm of Fiji (see methods for details). (C) Increased sensitivity of *syt1* compared to WT and *syt3* roots. (D) The cell viability of *syt1* roots is complemented by a SYT1-GFP fusion protein driven by the *SYT1* promoter. (E) *syt1/3* double mutant shows decreased cell viability compared to the single *syt1* mutant. (F) Cell viability of *syt1* roots is complemented by a SYT3-GFP fusion protein driven by the 35S promoter. Significant different values are indicated by different letters (P < 0.001; one-way ANOVA uncorrected Fisher’s LSD). Scale bars 50 µm. (G) Three-week-old WT, *syt1, syt3* and *syt1/3* Arabidopsis plants were grown under long-day photoperiod at 20°C and then cold-acclimated for 7 days at 4°C. After that, plants were exposed to different freezing temperatures for 6 h. Freezing tolerance was estimated as the percentage of plants surviving each specific temperature after 7 days of recovery under control conditions. Asterisks indicate statistical differences between mutant vs WT determined for each freezing temperature by one-way ANOVA and Tukey’s multiple comparison test (* p < 0.05, ** p < 0.005, **** p < 0.0001). Data represent mean values, error bars are SD, n _≥_ 30 plants per genotype for each freezing temperature.

Cell viability in roots of wild type (WT), *syt1* and *syt3* (Figure S1E) Arabidopsis seedlings was analyzed by confocal microscopy using fluorescein diacetate (FDA), a dye that only becomes fluorescent after its hydrolysis by esterases in living cells (Schapire et al., 2008). Thirty minutes of cold treatment (8°C) did not affect the cell survival of WT nor *syt3* roots, while it caused a dramatic decrease in cell viability of *syt1* roots (∼94%) (Figure 1C), quantified as the percentage of root area with visible FDA fluorescence above a threshold (see Methods). A genomic *SYT1* construct carrying *GFP* at the C-terminus (*syt1/SYT1:SYT1-GFP*) complemented the cell viability phenotype of *syt1* (Figure 1D) indicating that SYT1-GFP is functional. In order to investigate whether *SYT3* has a role in cell viability which might be masked by the highly expressed *SYT1*, we generated a *syt1/3* double mutant. Remarkably, after 10 min of cold treatment, an early time point at which no defects of cell viability are observed in *syt1*, around 80% of the *syt1/3* cells were already dead (Figure 1E). We then transformed Arabidopsis plants with genomic *SYT3* constructs fused to GFP at the C-terminus and driven by either *SYT3* promoter or the constitutive 35S promoter. Immunoblot analysis indicated a protein accumulation in the *35S:SYT3-GFP* line comparable to that of *SYT1:SYT1-GFP*, while we were not able to detect any signal in the *SYT3:SYT3-GFP* line (Figure S1F). Introducing the *35S:SYT3-GFP* construct in the *syt1* mutant background (*syt1*/*35S:SYT3-GFP)* complemented the *syt1* hypersensitive phenotype. Taken together, these results indicate that SYT1 and SYT3 can functionally complement each other when expressed at similar levels.

The *in silico* expression data also indicated that, in addition to cold, *SYT1* and particularly *SYT3* are regulated by different environmental stresses (Figure S1C and S1D), possibly reflecting a broad functionality of both proteins in abiotic stress tolerance. Therefore, we investigated the sensitivity of WT, *syt1, syt3*, and *syt1/3* plants to other abiotic stresses such as salt and osmotic stress. First, we analyzed root cell viability under salt stress of WT, *syt1, syt3*, and *syt1/3*. While *syt1* roots showed increased damage compared to WT roots, the sensitivity of *syt3* was very similar to WT. However, *syt1/3* roots showed significantly higher cellular damage than *syt1* under salt stress (Figure S1G). We next analyzed the responses of whole seedlings to osmotic stress by treating them with 20% of polyethylene glycol (PEG) and monitoring the cellular damage over time by measuring ion leakage. As shown in Figure S1H, we observed a similar damage in WT and *syt3* seedlings, and an increased damage in *syt1* that was further enhanced in *syt1/3*.

Cold acclimation is an adaptive response by which certain plants increase their freezing tolerance after being exposed for some days to low non-freezing temperatures (Thomashow, 1999) and previous reports have shown a role of *SYT1* in cold-acclimated freezing tolerance (Yamazaki et al., 2008). Therefore, we evaluated the involvement of *SYT1* and *SYT3* in cold-acclimated freezing tolerance in adult plants by measuring the survival rate of cold-acclimated (7 days, 4 °C) WT, *syt1, syt3* and *syt1/3* plants after exposure to freezing temperatures. As for other stresses, *syt3* plants exhibited a survival rate after freezing similar to WT (Figure 1G), *syt1* plants showed higher sensitivity than WT plants, while the *syt1/3* double mutant showed a freezing sensitive higher than *syt1* (Figure 1G). Taken together, our data indicate that *SYT1* and *SYT3* play a redundant role in the maintenance of PM stability in response to diverse abiotic stresses.

### Cold stress activates cellular damage responses in the *syt1/3* mutant

Cold acclimation involves an array of well-characterized transcriptomic modifications. We selected four marker genes of cold acclimation in Arabidopsis (Gilmour et al., 1988), *RCI2A, RAB18, COR78* and *COR27*, and analyzed their expression using RT-qPCR after 1, 3, and 7 days of exposure to 4 °C in WT and *syt1/3* plants. As shown in Figure S2A, no differences in the expression of these genes were found between WT and *syt1/3* plants, with all genes showing their maximum expression after 1 day of cold treatment. We next investigated the cold response of *CBF1* and *CBF3*, two Arabidopsis transcription factors that are critical for cold acclimation (Thomashow, 2010). As in our previous results, WT and *syt1/3* plants showed a similar induction of *CBF1* and *CBF3* after six hours at 4 °C (Figure S2B), suggesting that the *syt1/3* defects in cold acclimation are not due to an impaired transcriptional response. RNA-sequencing in WT and *syt1/3* plants at standard growth conditions and after 1 day at 4°C (the time point with the highest induction of the marker genes, see Table S1) revealed 1411 common cold up-regulated genes in both WT and *syt1/3*. Gene Ontology (GO) analysis of these up-regulated genes showed a significant enrichment in the categories “response to cold” and “response to temperature stimulus” (Table S2), indicating that the transcriptome of *syt1/3* is responding to cold.

Analysis of differentially expressed genes (DEG) in *syt1/3* relative to WT showed only 11 DEG between the two genotypes in standard growth conditions, with 8 genes being repressed and 3 induced in *syt1/3* relative to WT (Figure 2A). This result suggests that *SYT1* and *SYT3* have no obvious impact on the overall steady-state transcriptome of unstressed plants. Remarkably, cold treatment caused a significant impact on DEG with a total of 343 induced and 48 repressed genes in *syt1/3* relative to WT (Figure 2A, Table S1, Figure S2C). Functional enrichment of the 343 cold-induced genes in *syt1/3* relative to WT unveiled a significant over-representation of GO terms related to several responses to stress, both biotic and abiotic (Figure 2B, Table S3), while no GO terms were enriched in the 48 suppressed genes.

**Figure 2.**
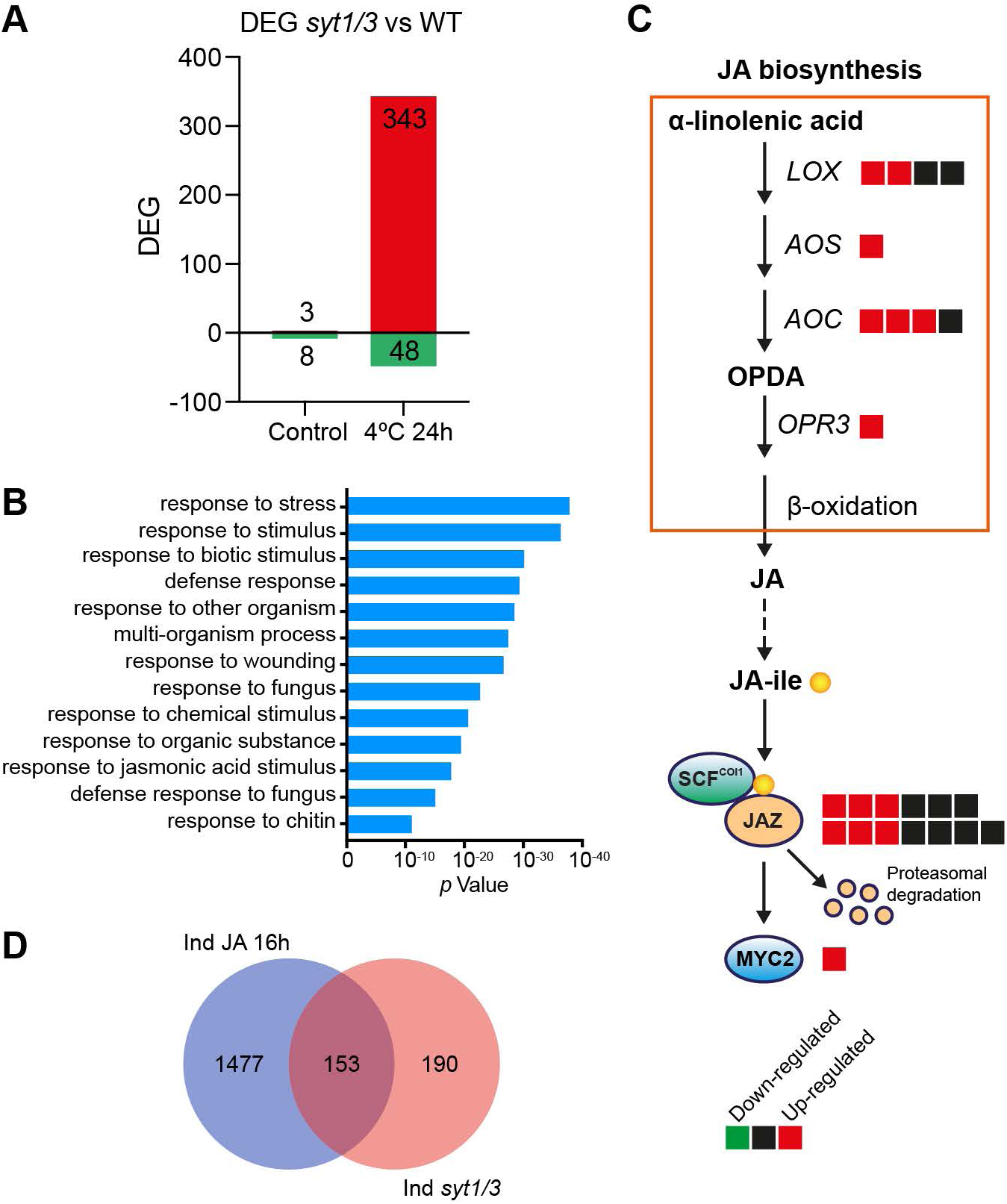
Cold stress activates cellular damage responses in the *syt1/3* mutant. (A) Differentially expressed genes (DEG) between WT and *syt1/3* in control conditions or after 24 h at 4°C treatment. Induced genes in *syt1/3* vs. WT are shown in red and repressed genes in *syt1/3* vs. WT are shown in green. (B) Gene Ontology terms (Biological Process assignments by TAIR/TIGR) over-represented among genes induced in *syt1/3* vs. WT. (C) Genes involved in the biosynthesis of jasmonic acid and transcriptional regulation of jasmonic acid responses. Squares indicated the number of genes identified at each step of the pathway. Red squares indicate genes that are induced in *syt1/3* vs. WT while black squares genes that are not induced. (D) Venn diagram showing overlap between genes induced in *syt1/3* vs. WT and genes induced after 16 h of JA treatment (Hickman et al., 2017).

Particularly striking was the over-representation of genes involved in the response to wounding and jasmonic acid (JA), a hormone involved in wound signaling (Sheard et al., 2010). The gene that showed the highest difference in expression (956 RPKM in *syt1/3* relative to WT) was lipoxygenase 2 (*LOX2*), a chloroplast enzyme required for wound-induced JA synthesis (Goossens et al., 2016). In addition to *LOX2*, genes involved in all steps of JA biosynthesis and transcriptional responses were also highly induced (Figure 2C).

Finally, we cross-referenced the list of cold-induced genes in *syt1/3* relative to WT with a published transcriptome of JA DEG (Hickman et al., 2017). Out of the 343 cold-induced genes in *syt1/3*, 153 (45 %) are also induced after 16 hr of JA (Figure 2D). These analyses reveal that while transcriptional responses to cold are similar in WT and *syt1/3* plants, there is a differential activation after cold treatment of wound and JA biosynthesis and signaling, suggesting that cold stress is perceived by *syt1/3* as a mechanical damage.

### SYT3 localizes at ER-PM CS

*SYT3* is reported as a pseudogene based on the presence of a stop codon in a cDNA sequence deposited in the Arabidopsis Information Resource (TAIR) Database (Yamazaki et al., 2008). The predicted protein lacks the C2 domains and is therefore expected to be non-functional. This observation is, however, in clear disagreement with the role we have identified in cell viability under abiotic stress (Figure 1E). To investigate this incongruity, we performed a detailed sequence analysis using available RNAseq databases and RT-PCR amplification followed by sequencing of the cDNAs fragments (Figure S3A) and identified a total of five splice variants (Figure S3B), including a transcript encoding a full SYT3 protein (*AT5G04220*.*1*), which contained similar domains to SYT1: a predicted TM region, an SMP domain and two C2 domains (Figure S3C).

The analogous modular structure of SYT1 and SYT3 and their redundant function in stress resistance suggest that, like SYT1, SYT3 is also an ER-PM CS tether. SYT3-GFP expressed in *Nicotiana benthamiana* leaves showed a reticulated beads-and-string pattern at the cortical plane of the cell (Figure 3A) and formed focused puncta around the cell periphery at the equatorial plane (Figure 3B), a localization pattern reminiscent of the ER-PM tethers SYT1-GFP and GFP-MAPPER (Figure S4A and S4B), an artificial ER-PM CS marker that localize at ER-PM CS (Lee et al., 2019) (Perez Sancho et al., 2015; Wang et al., 2014). Co-expression of binary combinations of differentially tagged constructs of SYT3, SYT1 and MAPPER (Figure 3C, Figure S4C, Figure S4D), produced a highly overlapping pattern. When we co-expressed SYT3-GFP with the ER localized proteins DGK2-mCherry (Angkawijaya et al., 2020) or FaFAH1-mCherry (Sánchez-Sevilla et al., 2014) we found that SYT3-GFP was mainly located in ER tubules and at the edges of ER sheets (Figure 3D, Figure S4E and Figure S4F). This localization is similar to that of Arabidopsis SYT1 (Ishikawa et al., 2018) and yeast Tcbs (Collado et al., 2019; Hoffmann et al., 2019), which have been reported to have important implications in the formation of the tubular ER network and in the formation of peaks of extreme curvature respectively.

**Figure 3.**
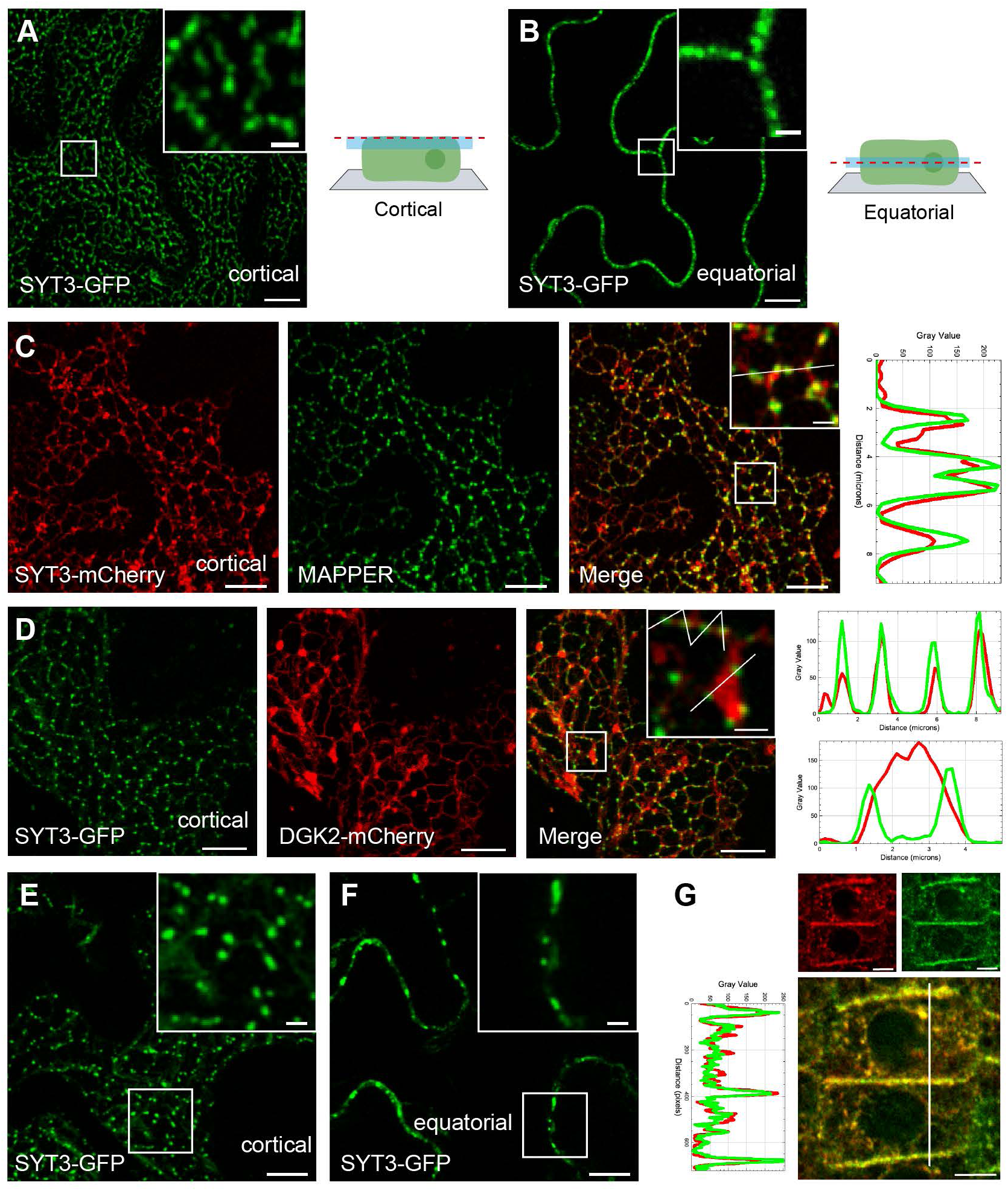
SYT3 localizes at ER-PM CS. (A-B) SYT3-GFP shows a beads-and-string pattern in *N. benthamiana* leaves. Confocal images of lower epidermis cells transiently expressing *SYT3-GFP* and schematic representations of the cell planes that are showed in the images, either at the cortical region (A) or at the equatorial plane (B). Boxed regions are magnified in the insets (close up). See also Figure S4A and Figure S4B. (C) SYT3-mCherry colocalizes with ER-PM CS marker. Confocal images of *N. benthamiana* leaves transiently co-expressing *SYT3-mCherry* and the ER-PM CS marker *MAPPER*, showing the cortical region as in (A). Images of the individual channels as well as the merged are shown. See also Figure S4C and Figure S4D. (D) SYT3-GFP localizes to ER tubules and the edges of ER sheets. Confocal images of the lower epidermis cells of *N. benthamiana* leaves transiently co-expressing *SYT3-GFP* with the ER marker *DGK2-mCherry*. Images show the cortical region of the cells as in (A). Images of the individual channels as well as the merged images are shown. See also Figure S4E, Figure S4F and Figure S4G. (E-F) SYT3-GFP shows a beads-and-string pattern in Arabidopsis. Confocal images showing the localization of SYT3-GFP either at the cortical region of the cells (E) or at the equatorial plane (F) in epidermal cells from the cotyledon of six-day-old Arabidopsis seedlings expressing *35S:SYT3-GFP* grown in half-strength MS agar solidified medium under long-day photoperiod and 23 °C. Boxed regions are magnified in the insets (close up). See also Figure S3D, Figure S3E and Figure S3F. (G) SYT3-GFP colocalizes with endogenous SYT1 in Arabidopsis roots. Five-day-old Arabidopsis seedlings expressing *35S:SYT3-GFP* grown in half-strength MS agar solidified medium under long-day photoperiod and 23 °C were fixed with 4 % PFA and permeabilized. The endogenous SYT1 protein was detected in the root tip with an anti-SYT1 primary antibody and a TRITC conjugated secondary antibody. Images of the individual channels as well as the merged image are shown. Scale bar 5 µm. See also Figure S3G. (A to F) Scale bars, 10 µm and scale bars for the close-up insets, 2 µm. (A to G) Cortical plane images are a maximum Z-projection of several planes from the cell surface until a plane where cells are close but still not touching the neighbors. Equatorial plane images are single plane images Intensity plots along the white lines in close up views are shown for each co-localization pattern.

In the stably transformed Arabidopsis *SYT3:SYT3-GFP* plants (Figure S1F), GFP signal was only observed in stomata and vascular tissues of primary roots (Figure S3D), indicating a low expression of SYT3-GFP under its own promoter. Consistently, a transgenic line with the reporter β*-glucuronidase* (*GUS*) gene under control of a 1,9 Kb *SYT3 cis*-regulatory region showed a similar expression pattern (Figure S3E). Due to this low signal, we further characterized the subcellular localization of SYT3 using the reported functional *35S:SYT3-GFP* Arabidopsis line (Figure 1E). Similar to SYT1-GFP (Figure S3F), SYT3-GFP showed a beads-and-string pattern at the cortical plane of the cell (Figure 3E) and focused puncta around the cell periphery at the equatorial plane (Figure 3F). In addition, using specific antibodies against SYT1 to immunolocalize endogenous SYT1 in the *SYT3:SYT3-GFP* line we obtained highly overlapping patterns in the roots (Figure 3G and Figure S3G) (Perez Sancho et al., 2015). Together, the experiments described above indicate that SYT3 localizes together with SYT1 at ER-PM CS.

### SYT1 and SYT3 tether the PM through interaction with PI4P

The expansion of ER-PM CS tethering is a defining property of ER-PM tethers when over-expressed (Eisenberg-Bord et al., 2016). Interestingly, in *N benthamiana* leaf epidermal cells where GFP-MAPPER was expressed alone, it showed a characteristic punctate pattern (Figure 4A, cell on the left and close up 1), but when it was co-expressed with SYT3-mCherry, MAPPER and SYT3 signals spread and colocalized throughout the entire cER (Figure 4A, cell on the right and close up 2), indicating the expansion of ER-PM CS. A similar effect was observed when GFP-MAPPER and SYT1-mCherry were co-expressed, occasionally causing the almost complete attachment of the ER to the PM (Figure S5A), a phenomenon that was associated to leaf necrosis (Figure S5B).

**Figure 4.**
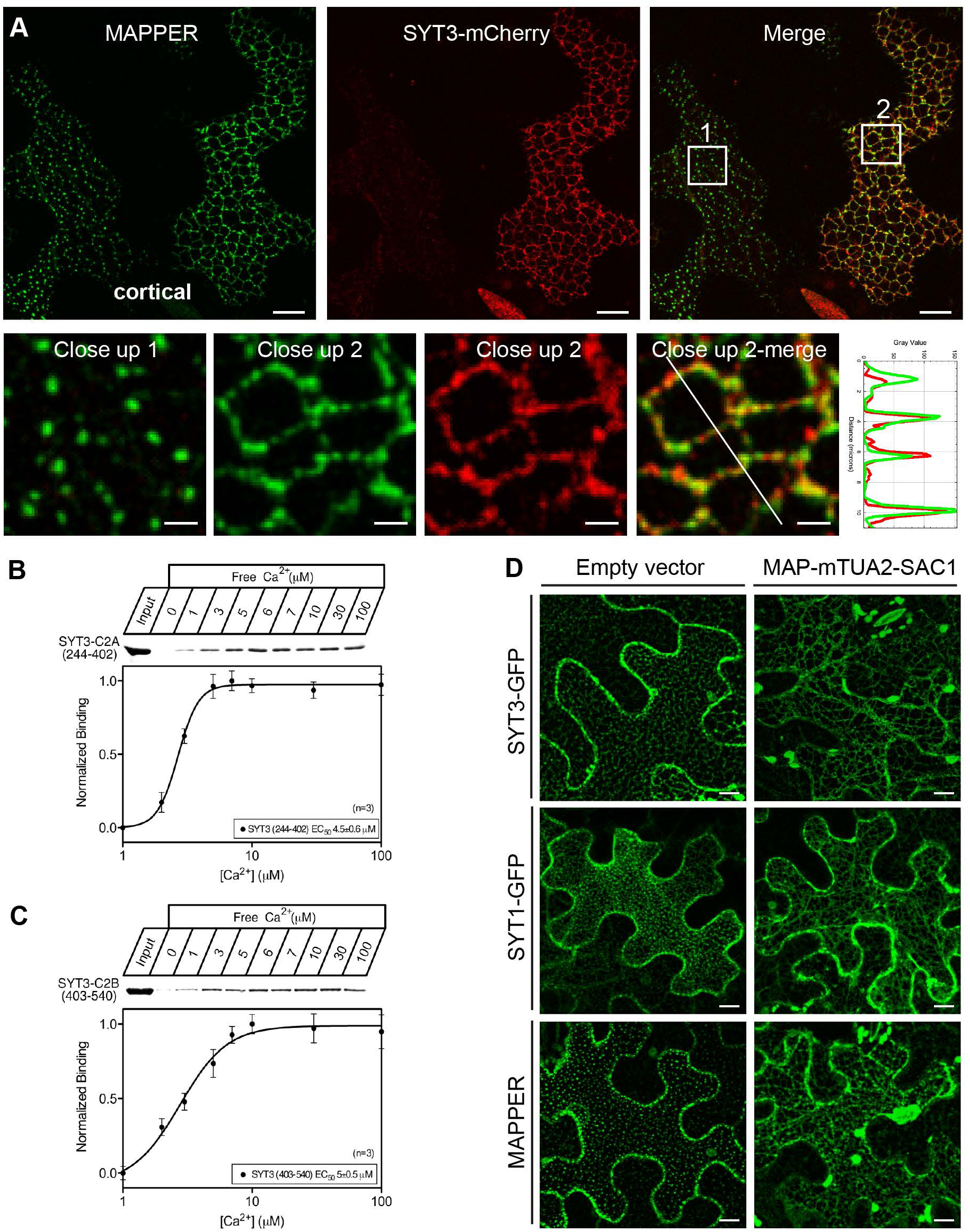
SYT1 and SYT3 tether the PM through interaction with PI4P. (A) *SYT3-mCherry* expression expands ER-PM CS in *N. benthamiana*. Confocal images of the cortical region of lower epidermis cells of *N. benthamiana* leaves transiently co-expressing *SYT3-mCherry* and the ER-PM marker *MAPPER*. Two cells are shown, one cell expressing only *MAPPER* (left cell, close up 1) and one co-expressing *SYT3-mCherry* and *MAPPER* (right cell, close up 2). Images of the individual channels as well as the merged images are shown. Boxed regions are magnified in the close ups. The intensity plot along the white line in close up 2 is shown. Scale bar, 10 µm and scale bar for close-up images, 2 µm. (B-C) Recombinant GST fusions of C2A domain (B) and C2B domain (C) of SYT3 containing the indicated residues were used in phospholipid binding assays. SYT3 domains were incubated with liposomes composed of 25% PS/75% PC in the indicated concentrations of free Ca^2+^ (clamped with Ca /EGTA buffers) in order to determine the half-maximal binding for free Ca^2+^. Liposomes were precipitated by centrifugation, and bound proteins were analyzed by SDS-PAGE. The EC_50_ for SYT3-C2A was 4,5 ± 0,6 µM and for SYT3 C2B was 5 ± 0,5 µM (paired *t* test, p < 0.05; n=4). (D) SYT3-GFP, SYT1-GFP and MAPPER localization on ER-PM CS depends on PM PI4P. Confocal images of the cortical region of lower epidermis cells of *N. benthamiana* leaves transiently co-expressing *SYT3-GFP* (left row), *SYT1-GFP* (central row) or *MAPPER* (right row) either with an empty vector (top panels) or with the PM bound PI4P phosphatase MAP-mTU2-SAC1. Scale bars 10 µm.

Analogously to SYT1 (Perez Sancho et al., 2015), SYT3 is expected to be anchored to the ER and to bind in *trans* to the PM via its C2 domains (Figure S3C). Earlier work has shown that the C2A domain of SYT1 binds artificial liposomes containing phosphatidylserine (PS) and PC in a Ca^2+^-dependent manner, while binding of SYT1-C2B is independent of Ca^2+^ (Schapire et al., 2008). Calcium binding sites in C2 domains are characterized by acidic residues (Asp/Glu) confined in conserved loops that play crucial roles in coordinating Ca^2+^ ions (Fernández-Chacón et al., 2001). Sequence analysis of the C2 domains of SYT1, SYT3 and E-Syt1 predicted the presence of conserved Ca^2+^-binding sites in the C2A domains of SYT1 and SYT3 and in the C2A and C2C of E-Syt1 (Figure S5C), as previously reported (Giordano et al., 2013). Additionally, 3-D modeling of the independent C2A and C2B domains of SYT3 identified a Ca^2+^-binding pocket in the C2A of SYT3 (Figure S5D). Nevertheless, since Ca^2+^ and phospholipid binding properties of C2 domains cannot be reliably predicted from sequence analysis (Dai et al., 2004), we investigated the Ca^2+^-dependent phospholipid binding properties of SYT3 C2A and C2B domains. As shown in Figure 4B, the purified SYT3-C2A domain fused to glutathione-S-transferase (SYT3-C2A-GST) bound negatively charged liposomes (25% PS/75% PC) in a Ca^2+^-dependent manner with an estimated half-maximal Ca^2+^ binding value of 4.5 ± 0.6 µM free Ca^2+^, which is slightly smaller than that of SYT1-C2A (6 ± 0.6 µM free Ca^2+^) (Schapire et al., 2008) and very similar to the C2A of rat synaptotagmin 1 (4.1 ± 0.3 µM free Ca^2+^) (Fernández-Chacón et al., 2001; Schapire et al., 2008). However, SYT3-C2B-GST showed some Ca^2+^-independent binding to these liposomes and its binding increased in the presence of Ca^2+^ (EC = 5.0 ± 0.5 µM free Ca^2+^) (Figure 4C). These results indicate that the C2B domain of SYT3 might contain additional Ca^2+^ binding sites that are different from the canonical ones. Indeed, non-canonical Ca^2+^ binding sites have been reported for the C2AB fragment of human E-Syt2 (Xu et al., 2014). In both cases, the affinity of Ca^2+^ binding for C2A and C2B domains are in the low µM range, within the concentration ranges considered physiological in plants and mammals (Bian et al., 2018; H. Knight et al., 1997).

C2 domains, in addition to PS, can bind other negatively charged phospholipids (Schapire et al., 2008) (Perez Sancho et al., 2015). Because PI4P is the main determinant generating negative charges at the inner surface of the PM (Simon et al., 2016), we investigated the role of PI4P in the localization of SYT3-GFP and SYT1-GFP at ER-PM CS *in vivo*. In order to do so, SYT3-GFP and SYT1-GFP were co-expressed in *N. benthamiana* with the PM anchored MAP-mTU2-SAC1 phosphatase (Simon et al., 2016), which depletes PI4P from the PM (Simon et al., 2016). Additionally, we used the artificial tether GFP-MAPPER, which binds the PM through the interaction of a polybasic domain with negatively charged phospholipids (Chang et al., 2013). Expression of individual SYT1-GFP, SYT3-GFP, and GFP-MAPPER resulted in a typical cortical ER-PM CS localization (Figure 4D, left panels). Nonetheless, co-expression of the same constructs with MAPmTU2-SAC1 caused the redistribution of the GFP signals throughout the ER (Figure 4D, right panels). In an alternative approach, we decreased the content of PI4P at the PM in Arabidopsis roots by the application of phenylarsine oxide (PAO), an inhibitor of the phosphatidylinositol-4-kinase (PI4K) (Simon et al., 2016). PAO treatment in a transgenic line expressing SYT1-GFP and the PI4P biosensor 1xPH^FAPP1^ triggered the dissociation of the biosensor from the PM into the cytosol (Figure S5E, left panels) and caused a marked increase in perinuclear staining of SYT1-GFP (Figure S5E, central and right -detail-panels), indicating a SYT1-GFP relocalization from ER-PM CS to a broader distribution in the ER. These results are consistent with a tethering role for SYT1 and SYT3 in which their N-terminal TM domain anchors them to the ER and their C-terminal C2 domains bind *in trans* to the negatively charged PM.

### SYT1 and SYT3 show cold-dependent dynamics at ER-PM CS

SYT1 was originally identified as a PM protein that accumulates in the PM fractions in response to cold stress (Kawamura and Uemura, 2003). However, using specific SYT1 antibodies (Perez Sancho et al., 2015) we did not detect increased accumulation of total SYT1 protein after 1, 3 or 7 days of cold exposure in WT or in the *syt3* mutant (Figure 5A). This indicates that the total amount of SYT1 protein is neither altered by cold treatment nor by the absence of SYT3. Our results open the possibility that SYT1 accumulation at the PM fraction is the consequence of a higher concentration at ER-PM CS (Kawamura and Uemura, 2003). *SYT3* expression is induced by several stresses, including cold (Figure 1B, S1C and S1D), therefore we investigate the level of SYT3-GFP protein after cold in the *SYT3:SYT3-GFP* line using immunoblot. However, the amount of SYT3-GFP remained below the detection level (Figure 5B and Figure S1F). Therefore, SYT3-GFP was concentrated using GFP-trap columns, and contrary to SYT1, cold treatment caused a ∼6-fold induction of SYT3-GFP (Figure 5B).

**Figure 5.**
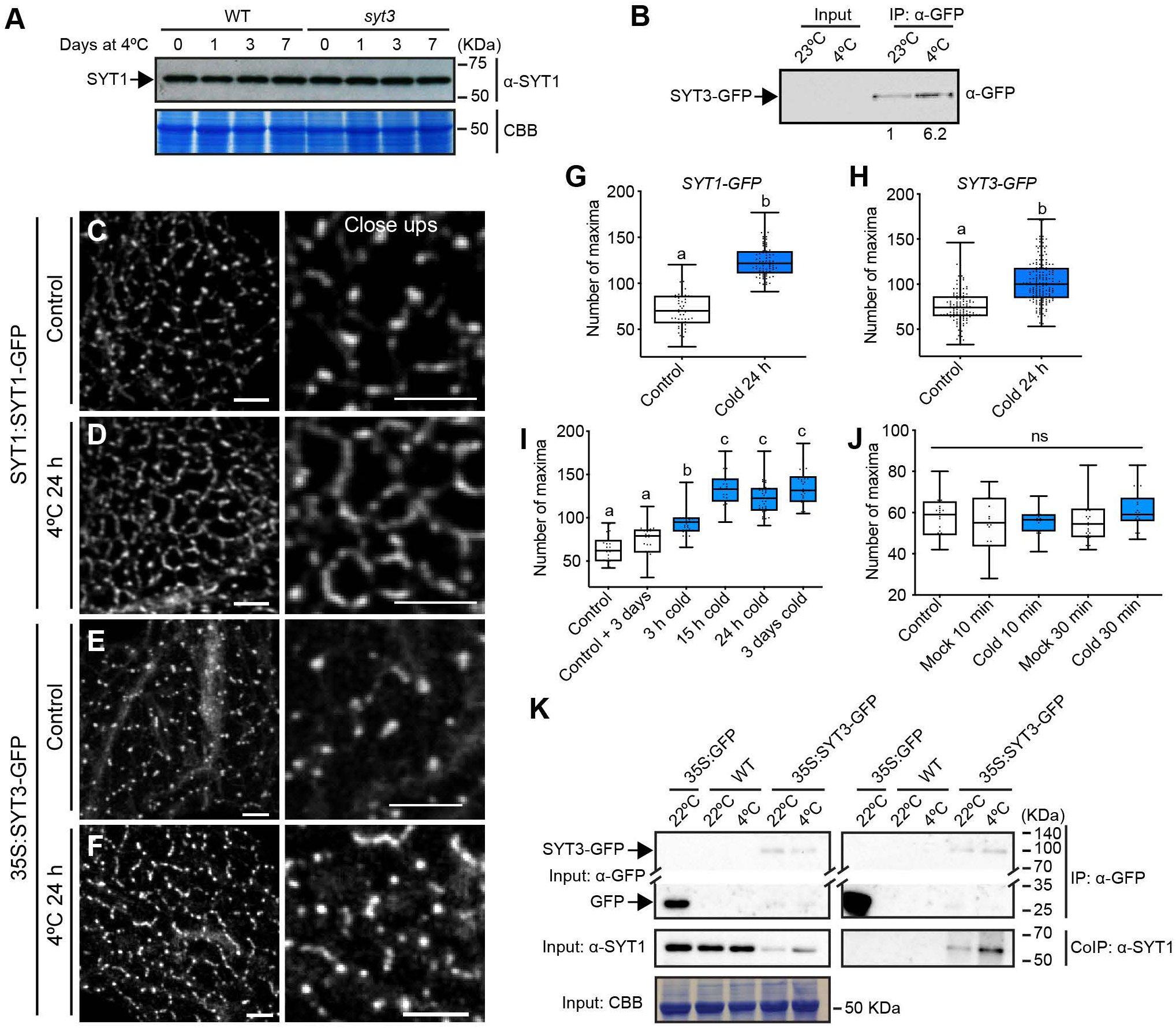
SYT1 and SYT3 show cold-dependent dynamics at ER-PM CS. Total SYT1 protein level does not change after cold treatment. Arabidopsis WT and *syt3* plants were grown for 3 weeks on soil under long-day photoperiod at 20°C and then transferred to 4°C for 1, 3 or 7 days. Proteins were extracted from shoots and detected by immunoblot with anti-SYT1 antibody. Equal loading was confirmed by Coomassie blue staining (CBB). Experiment was repeated three times with similar results. SYT3-GFP protein accumulation is induced by cold. Arabidopsis seedlings expressing *SYT3:SYT3-GFP* were grown under long-day photoperiod and 23°C for 7 days and then transferred to 4°C for 24 h or kept at 23°C. Proteins from whole seedlings were detected by immunoblot with anti-GFP antibody. No signal was detected in the crude extracts (INPUT). SYT3-GFP was concentrated using GFP-Trap beads (GFP-IP), detected by immunoblot with anti-GFP antibody and quantified using FIJI. (C-F) Representative confocal images showing SYT1-GFP (C and D) and SYT3-GFP (E and F) dynamics with cold. Arabidopsis seedlings expressing *SYT1:SYT1-GFP* or *35S:SYT3-GFP* were grown under long-day photoperiod and 23°C for 7 days. Cold treatment plates were transferred to 4°C for 24 h (D and F) while control plates kept growing in control conditions (C and E). Mounting of the cold-treated seedlings in the coverlids was done under cold conditions and with pre-chilled water. Images show a maximum Z-projection of the cortical region of the epidermal cells from the cotyledon. Close ups are shown for detailed view. Scale bars 10 µm and 5 µm for the close-ups. (G-H) Quantification of ER-PM CS labeled by SYT1-GFP (G) or SYT3-GFP (H) in cotyledons of 7-day-old Arabidopsis epidermal cells in control conditions or after 24 h at 4°C. Contacts sites were identified as intensity maxima in the cortical region using FIJI (see methods for details). Dots represent individual measurements from independent ROI from at least 5 cotyledons. Box plots display the first and third quartiles, split by the median; whiskers extend to include the maximum and minimum values. Different lowercase letters indicate significant differences. Data were analyzed with one-way ANOVA and Tukeys multiple comparison test; p < 0.05. Quantification of the amount of SYT1-GFP labeled ER-PM CS in the epidermal cells of the cotyledon of 7-day-old Arabidopsis seedlings in control conditions or after 3 h, 15 h, 24 h and 3 days at 4°C. “Control + 3 days” is included to show that differences in the amount of CS are not due to cotyledon age. Contacts sites were identified as intensity maxima in the cortical region using FIJI (see methods for details). Dots represent individual measurements from independent ROI from at least 5 cotyledons. Statistical analysis as in panels G-H. Quantification of SYT1-GFP labeled contact sites in the epidermal cells of the cotyledon of 7-day-old Arabidopsis seedlings in mock conditions or after 10 min and 30 min of cold treatment. Arabidopsis seedlings expressing *SYT1:SYT1-GFP* were grown under long-day photoperiod and 23°C for 7 days. Seedlings for cold treatment were covered with liquid half-strength MS pre-chilled at 4°C and the plates were transferred to a box with ice for the specified times. Mounting of the seedlings in the cover-slides was done under cold conditions and with pre-chilled water. Mock seedlings were covered with liquid half-strength MS at 23°C for the specified times. Contacts sites and statistical analysis as were quantified as described in panels G to J. SYT1 co-immunoprecipitates with SYT3-GFP. Thirteen-day-old Arabidopsis WT, and expressing *35S:GFP* and *35S:SYT3-GFP* were subjected to 23°C and 4°C during 3 days. Total (input), immunoprecipitated (IP) and Co-Immunoprecipitated (CoIP) proteins were analyzed by immunoblotting. Equal loading in the input was confirmed by Coomassie blue staining (CBB). Free GFP was used as a negative control for Co-IP. SYT3-GFP and free GFP were detected with anti-GFP antibody and SYT1 was detected with anti-SYT1 antibody. Anti-GFP (input and IP) where crop and the top and bottom part of the blots that show the SYT3-GFP and free GFP protein, respectively, are shown.

Next, we investigated the dynamics under cold stress at the cellular level of SYT1-GFP using the *SYT1:SYT1-GFP* complementing line, and SYT3-GFP using the *35S:SYT3-GFP* complementing line (due the low expression of *SYT3* under its endogenous promoter). In control conditions, SYT1-GFP and SYT3-GFP were observed at cER, showing the characteristic discrete punctate pattern (Figure 5C, 5E). Although cold does not affect the total amount of endogenous SYT1 (Figure 5A) and SYT3-GFP (which is driven by the constitutive 35S promoter) proteins, we observed an increase of SYT1-GFP and SYT3-GFP signals at the cortical plane after 24 hr of cold treatment (Figure 5D, 5F). Image analysis (see Methods) confirmed significant changes in the localization of SYT1-GFP and SYT3-GFP after cold treatment (Figure 5G, 5H). These results are indicative of a recruitment of SYT1 and SYT3 to the ER-PM CS. Quantification of the distribution of SYT1-GFP at 3 hr, 15 hr, 24 hr and 3 days after cold treatment, indicated an increased re-localization at 3 hr that remained up to 3 days of cold treatment (Figure 5I). Surprisingly, 10 and 30 min of cold treatment did not produce any significant changes in the localization of SYT1 (Figure 5J). This indicate that the relocalization of plant SYTs at ER-PM CS after cold is slow and persistent, in contrast to the Ca^2+^-dependent recruitment to ER-PM CS of E-Syt1 that is very fast (within seconds/minutes) and transient (Giordano et al., 2013; Saheki et al., 2016). Thus, our results reveal striking differences in the dynamics between animal E-Syts and plant SYTs.

E-Syts form homo- and hetero-dimers and in fact, these interactions affect their respective localization and Ca^2+^-dependent lipid binding (Giordano et al., 2013). Arabidopsis SYT1 forms homodimers as well (Lee et al., 2020). To assess formation of heterodimers between SYT1 and SYT3, *35S:SYT3-GFP* plants in WT background were used and analyzed for their interaction using co-immunoprecipitation (Co-IP) assays in control conditions (22 °C) or after 3 days of cold treatment. In both conditions, SYT3-GFP Co-IP endogenous SYT1 proteins indicating an *in vivo* association (Figure 5K).

### SYT1 and SYT3 maintain DAG homeostasis at the PM upon cold

Human E-Syts contain a SMP domain with a hydrophobic groove that harbors glycerolipids (Schauder et al., 2014). Therefore, to evaluate a possible role of SYT1 and SYT3 proteins in glycerolipid homeostasis, we analyzed the leaf glycerolipidome in control conditions and after 7 days at 4°C of WT, *syt1, syt3* and *syt1/3* plants. Using ultra-performance liquid chromatography coupled to Fourier-transform mass spectrometry (UPLC-FT-MS)-based high-resolution lipidome analyses we profiled 181 molecular species from nine glycerolipid classes (Table S4). In WT plants, cold stress caused an increase in the amount of unsaturation in most lipid classes (Figure S6A), a decrease of monogalactosyldiacylglycerols (MGDG) and the accumulation of unsaturated triacylglycerols (TAG), which is consistent with previous reports (Degenkolbe et al., 2012; Tarazona et al., 2015). As shown in Figure 6A, the total glycerolipid composition of *syt1, syt3* and *syt1/3* were similar to that of WT plants in both control conditions and after a cold treatment, ruling out a role for SYT1 and SYT3 proteins in the substantial lipid remodeling that takes place after cold stress.

**Figure 6.**
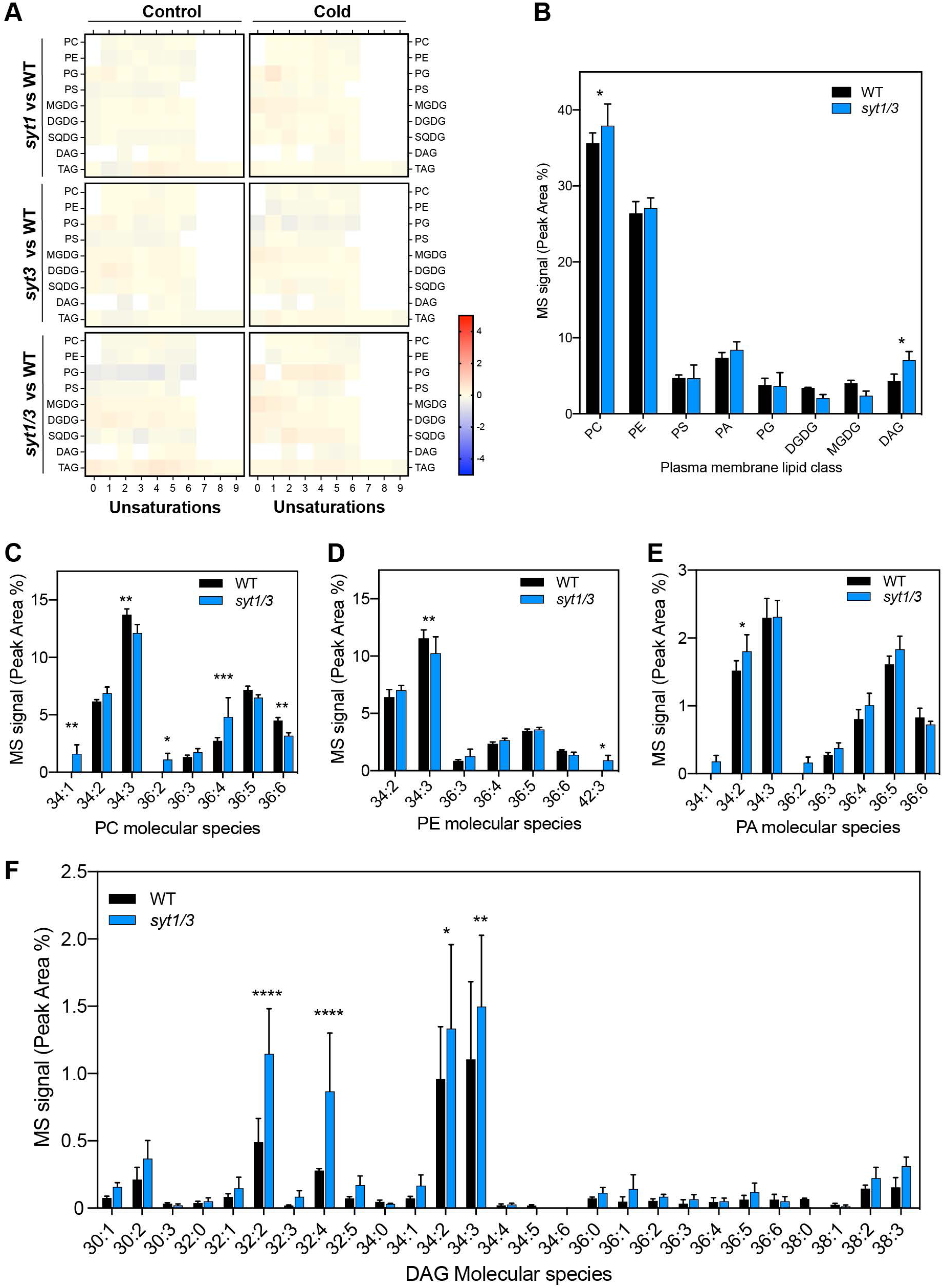
SYT1 and SYT3 maintain DAG homeostasis at the PM. Heatmap panels representing the fold change in the peak area of each glycerolipid class and unsaturation grade of the acyl chains when comparing WT against *syt1, syt3* or *syt1/3*. Four-week-old rosettes were analyzed after growing under control conditions and after a 7 days cold treatment (4°C). See also Figure S6A. Color scale code expresses fold changes of the corresponding glycerolipids (n = 3). ESI-MS/MS analysis of the molecular species of PM glycerolipids from 4-week-old WT and s*yt1/3* rosettes grown at 23°C followed by 3 days of cold treatment (4°C). Plasma membrane samples were purified by two phase partitioning protocol and lipids were extracted following as described in Methods (mean with SD, n=3). See also Figure S6B. (C-F) Distribution of the identified PC (C), PE (D), PA (E) and DAG (F) molecular species in PM of WT and *syt1/3*. Acyl chains are expressed as number of acyl carbons: number of acyl double bonds. Data of lipids in plasma membrane are represented as mean with standard deviation (n=3). Asterisks indicate statistically significant differences between PM in WT and *syt1/3* plants as determined by the Fisher LSD test; p < 0,0001(****); p < 0,0002(***); p < 0,0021 (**); p < 0,0332 (*). Abbreviations: PA, phosphatidic acid; PC, phosphatidylcholine; PE, phosphatidylethanolamine; PG, phosphatidylglycerol; PS, phosphatidylserine; MGDG, monogalactosyldiacylglycerol; DGDG, digalactosyldiacylglycerol; SQDG, sulfoquinovosyldiacylglycerol; DAG, diacylglycerol; TAG, triacylglycerol.

**Figure 7.**
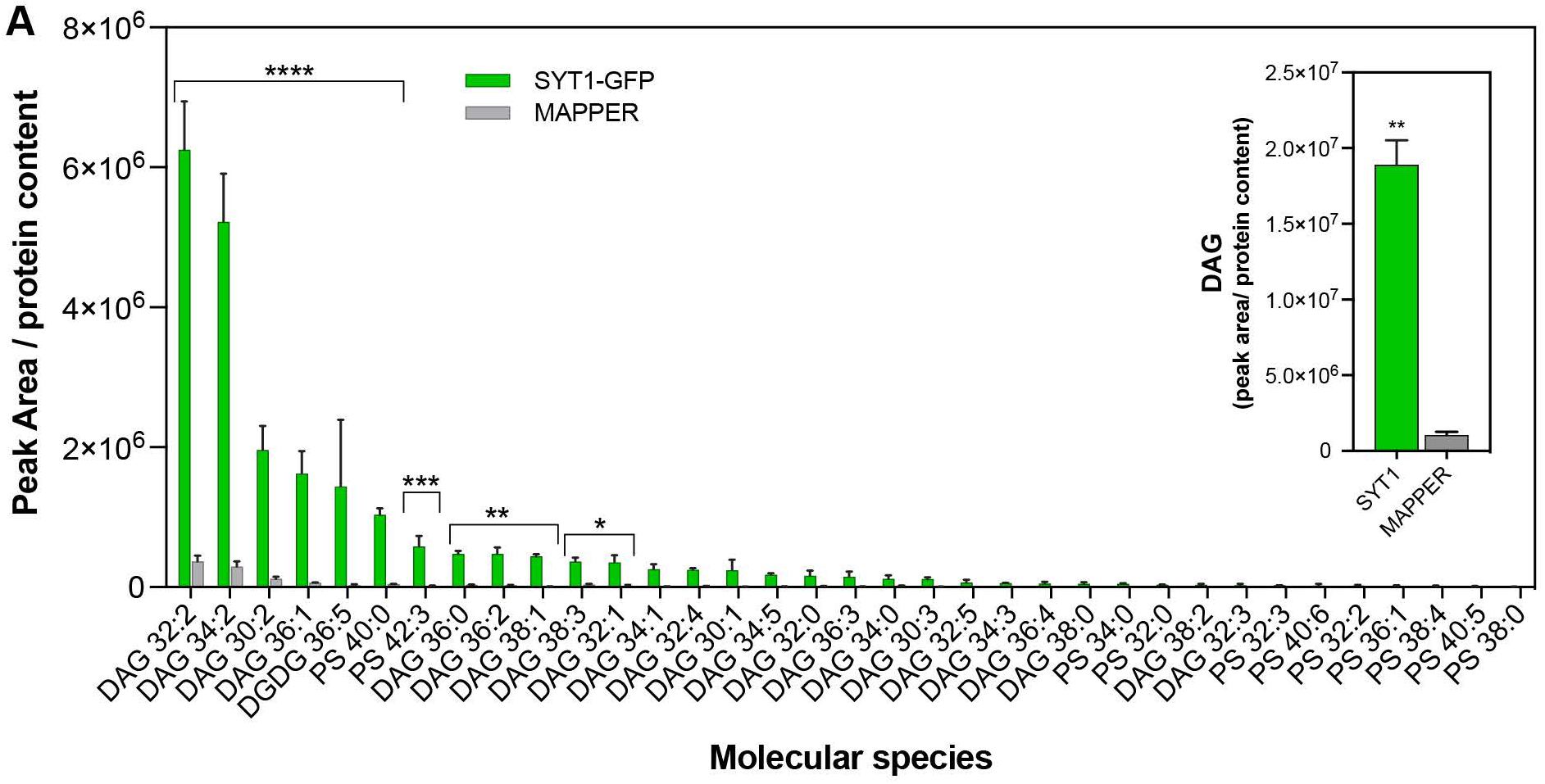
SMP domain of SYT1 preferentially binds to DAGs *in vivo*. (A) Glycerolipid molecular species determined by ESI-MS/MS analysis and identified in the immunoprecipitated SYT1-GFP and MAPPER proteins from 4-week-old Arabidopsis plants grown at 23°C followed by a 3 days cold treatment (4°C). Leaves of Arabidopsis plants expressing *SYT1-GFP* or *MAPPER* proteins were homogenized, and proteins were affinity purified using beads coated with anti-GFP antibody. Lipids were extracted using Bligh and Dyer protocol. Acyl chains are expressed as [number of acyl carbons: number of acyl double bonds]. The inset graph shows the comparison of total DAG in SYT1-GFP and MAPPER immunoprecipitated proteins. Verification that SYT1-GFP did not co-IP with MAPPER is in Figure S6A. Lipid species were expressed as mean peak area. Error bars indicate SD. Asterisks indicate statistically significant differences between SYT1-GFP and MAPPER lipids as determined by the Fisher LSD test; p < 0,0001 (****); p < 0,0002 (***); p < 0,0021 (**); p < 0,0332 (*).

The localization and the increased accumulation of SYT1 and SYT3 at ER-PM CS point towards a lipid redistribution between ER and PM more than to a general lipid remodeling. Thus, we investigated lipid changes specifically in the PM of WT and *syt1/3* plants after three days of cold treatment, a time point in which SYT1 showed a strong re-localization at ER-PM CS. To achieve a good resolution, we utilized a two-phase partitioning protocol to obtain enriched PM fractions minimizing ER contaminations. Immunoblot analysis (Figure S6B) showed that the PM fraction was highly enriched in AHA3, a H^+^-ATPase localized at the PM, while the luminal binding protein BIP (a common ER marker covering BIP1, BIP2 and BIP3), and SYT1 were depleted. In contrast, BIP and SYT1 were enriched in endomembrane fractions (EM), while AHA3 was diminished. This analysis confirmed the purity of the PM fractions and corroborated the ER anchoring of SYT1 via its TM domain. Electrospray ionization tandem mass spectrometry (ESI-MS/MS) of the PM fractions allowed the quantification of 78 glycerolipids species belonging to nine lipid classes. The low amount of chloroplastic MGDG and digalactosildiacylglycerol (DGDG) lipids in the PM samples further supported the purity of the isolated membranes (Table S5). The analysis showed no significant changes in the composition of major lipids such as PE, PS, PG and PA between WT and *syt1/3* PM while significant differences were only found in PC and DAG (Figure 6B-E). Examination of PC species revealed a complex pattern of changes in *syt1/3* relative to WT (Figure 6C). Remarkably, most DAG molecular species displayed a tendency to accumulate in *syt1/3* PM (Figure 6F), with DAG32:2 and DAG32:4 being the most enriched ones.

### SYT1 preferentially bind to DAGs *in vivo*

SMP domain of E-Syts forms a hydrophobic groove that can transfer glycerolipids between membranes, but the nature of the transported glycerolipids and their directionality is not unequivocally resolved (Schauder et al., 2014). Therefore, we sought to identify glycerolipids bound to the SYT1 SMP domain *in vivo* by analyzing the glycerolipids that co-immunoprecipitated with the functional SYT1-GFP (*syt1/SYT1:SYT1-GFP* Arabidopsis line). The artificial ER-PM tether GFP-MAPPER was used as negative control because it co-localizes with SYT1 at ER-PM CS but does not contain a SMP domain. We first established that GFP-MAPPER does not co-immunoprecipitate SYT1 (Figure S6A), as this would render lipids in the GFP-MAPPER sample that would be actually bound to the co-purified SYT1. Then, SYT1-GFP and MAPPER where purified using CHAPS as detergent, which has been reported to not displace lipids bound to the SMP domain of Mdm12 (AhYoung et al., 2015). The lipids bound to the co-immunoprecipitated proteins where subsequently extracted and analyzed by electro spray ionization tandem mass spectrometry (ESI-MS/MS) (Table S6). Surprisingly, the analysis did not detect the presence of species of highly abundant lipid classes such as PC, PE, or phosphatidylglycerol (PG). Remarkably, out of the 35 lipid species identified in SYT1-GFP samples, 24 were DAGs (Figure 6C). The two most abundant DAG species enriched in SYT1-GFP were DAG32:2 and DAG34:2. In addition to DAGs, a single species of digalactosildiacylglycerol (DGDG36:5), PS40:0 and minor amounts of some PS species were also detected in SYT1-GFP. These results are consistent with the finding that most lipids bound to SYT1-GFP are molecular species of DAG and support a role for SYT1 and SYT3 in transporting DAGs from the PM to the ER.

## DISCUSSION

Our results provide direct evidences that SYT1 and SYT3 are implicated in the clearance of PM DAG that is produced during stress. The lack of these proteins in *syt1/3* plants results in the accumulation of DAG at the PM, that likely causes the defective PM integrity exhibited in response to multiple stresses and explains the induction of genes involved in wounding in response to cold. SYT1 and SYT3 localization at ER-PM CS is directed by their TM anchor to the ER and the binding of their C2 domains in *trans* to PM PI4P and follows and stress-dependent dynamics. Thus, the efficiency of DAG removal is likely enhanced by the SYT1 and SYT3 accumulation at ER-PM CS after abiotic stress.

### SYT1 and SYT3 function as dynamic ER-PM CS tethers in response to abiotic stress

SYT1 and SYT3 proteins function as tethers in the formation of ER-PM CS. This localization is dependent on their ER anchoring by their N-terminal transmembrane domain and the interaction of their C2 domains with PI4P at the PM. Consistently, SYT1 purifies within the ER fraction and the depletion of PI4P in the cytosolic leaflet of the PM by MAP-SAC1 triggers the loss of SYT1-GFP and SYT3-GFP focal localization at ER-PM CS, resulting in a distribution of these proteins throughout the ER. As it is anticipated for proteins that function as ER-PM tethers (Eisenberg-Bord et al., 2016), overexpression of SYT1 and SYT3 causes a drastic expansion of ER-PM CS that can eventually results in cell death. Interestingly, SYT3 shows an enrichment in ER domains that display high curvature, like tubules and sheet edges (Figure 3D and S4E, S4F, S4G). This localization to high ER curvature has been previously reported for SYT1 (Ishikawa et al., 2018) and Tcb proteins (Collado et al., 2019; Hoffmann et al., 2019), which may have important functional implications in lipid transport as will be later discussed. Cold stress increases the ER-PM connectivity by promoting the cortical expansion of ER-PM CS containing SYT1 and SYT3 without changes in the total amount of SYT1 (Figure 4A) nor SYT3, which is driven by the constitutive 35S promoter. A similar increase of ER-PM CS marked by SYT1 has been previously reported by salt stress (Lee et al., 2019), indicating that SYT1 (and likely SYT3) dynamics are not specific to cold and establishing SYT1 and SYT3 as general abiotic stress-regulated tethers between the ER and the PM.

An obvious question is what the molecular events that trigger SYT1 and SYT3 relocalization and ER-PM CS expansion are. Based on previous data from E-Syt1, one evident candidate that triggers SYT1 and SYT3 stress-dependent dynamics is cytosolic Ca^2+^. First, it is known that cytosolic Ca^2+^ increase as a response to multiple abiotic stresses (H. Knight et al., 1997; M. R. Knight et al., 1991). Second, C2 domains of SYT1 (Schapire et al., 2008) and SYT3 (this work) show phospholipid biding that are dependent on micromolar increase of Ca^2+^. However, the Arabidopsis temporal dynamics of SYT1 and SYT3 makes it difficult to reconcile a Ca^2+^ increase as being the molecular trigger for this re-localization. Changes in SYT1 localization are observed after ∼3 hours of cold and remained even after 3 days of treatment while cold stress is known to cause very fast and transient increases in cytosolic Ca^2+^ (H. Knight and M. R. Knight, 2000). Although we cannot rule out that the cold ER-PM CS expansion is a downstream response to the transient Ca^2+^, one obvious candidate for SYT1 and SYT3 dynamics is PI(4,5)P_2_. This lipid species is induced by different stresses at the PM (Heilmann, 2016) and salt stress causes a similar SYT1 relocalization at ER-PM CS that is associated with changes in the content of PI(4,5)P_2_ (Lee et al., 2019).

### A role for SYT1 and SYT3 for diacylglycerol homeostasis during abiotic stress

SYT1 is not required for the expansion of ER-PM CS after stress, suggesting additional functions for SYT1 other than ER-PM CS tethering (Lee et al., 2019). Clearly, a defective lipid transport associated to their SMP domains could be the cause underlying the *syt1/3* stress-dependent phenotypes. E-Syts, can promiscuously transfer glycerolipids between bilayers, including DAG (Bian et al., 2018; Saheki et al., 2016; H. Yu et al., 2016). Although a first metabolomic analysis did not reveal differences in the composition of glycerolipid species of total lipids, PM fractions from cold treated *syt1/3* plants showed a significant accumulation of DAG compared to the WT. Importantly, most DAG species were increased while significant changes were found in DAG32:2, DAG32:4, DAG34:2 and DAG34:3. Moreover, immunoprecipitation of SYT1-GFP followed by lipidomic analysis revealed the presence of many DAG species, suggesting a lack of specificity with respect to any particular DAG species.

Upon stress, PLCs and NPCs which hydrolyze PIPs and membrane structural phospholipids such as PC or PE respectively, can be activated triggering an increase of DAG at the PM. Although both the NPCs and PLCs can induce DAG formation, the DAG molecular species they produce are different due to the substrates they hydrolyze. While DAG produced via the PLC pathway is mainly composed of C32 and C34 species (König et al., 2008), NPC generates DAG species with longer (C34 and C36) and more unsaturated fatty acids (Peters et al., 2010). When analyzing the PM of *syt1/3*, we did not detect any significant changes in DAG molecular species with 36 carbons, and the biggest changes were found in shorter DAG with 32 carbons. These results suggest that DAG species in *syt1/3* PM are likely produced due to the activation of PLCs.

DAG molecules are conical in shape within the membrane and thus packs poorly in planar bilayers and generate areas of negative curvature leading to a disruption of the lamellar phase of lipid membranes (Campomanes et al., 2019). Consequently, it is predicted that an uncontrolled accumulation of DAG caused by mutations in *SYT1* and *SYT3* will lead to a severe loss of plasma membrane integrity, highlighting the importance of DAG removal during stress episodes. Interestingly, although no significant alterations in steady-state PM glycerolipids were detected in genome-edited cells lacking all E-Syts, a transient accumulation of DAG after PLC activation was observed, supporting their role in DAG transport (Saheki et al., 2016). This function was further reinforced from the analysis of pancreatic islets, where a decreased expression of *E-Syt1* resulted in prolonged accumulation of DAG in the PM causing an increase in insulin secretion (Xie et al., 2019).

Recently, it has been shown that Tcbs create peaks of extreme curvature on the cER and these peaks are important to maintain PM integrity under stress, likely by facilitating lipid transport (Collado et al., 2019). Like Tcbs, SYT1 and SYT3 are localized in regions of the ER that display high curvature (Figure 3D, Figure S4E and Figure S4F) (Ishikawa et al., 2018). Accordingly, it is conceivable that Tcbs, like SYTs and E-Syts, may also operate in the regulation of DAG levels at the PM and their specific localization play a role in the mechanism of lipid transfer. In pancreatic β cells, DAG formation occurs in microdomains that coincide with sites of insulin granule exocytosis and are accompanied by recruitment of E-Syt1 to the same sites. It is tempting to speculate that the formation of DAG by PLC in plants might occur in specific ER-PM CS microdomains that overlap with the localization of SYT1 and SYT3, increasing the efficiency of transfer.

The large evolutionary distance between eukaryotes such yeast, mammals and plants have resulted in the generation of independent mechanisms to deal with developmental and environmental challenges. However, they all have in common the presence of ER-PM CS that play important metabolic functions and express a class of closely related proteins. Mammal E-Syts, yeast Tcbs and plant SYTs localized in these microdomains, functioning as tethers that connect the ER with the PM. Still, whether or not these proteins have common roles in lipid transfer through their SMP domains remains elusive. This report provides novel insights into the role of the SMP domain containing proteins SYT1 and SYT3 that probably can be translated to yeast and mammals.

In summary, our studies have revealed a role for ER-PM CS SYT1 and SYT3 proteins in the regulation of PM DAG homeostasis under abiotic stress. We propose a model in which activation of PLC induces rapid and transient formation of DAG during stress episodes. Because of their structural characteristics, DAG does not form lipid bilayers and therefore must be actively removed. Part of this pool of DAG is transformed into PA by action of DAG kinases, while the remaining DAG is efficiently transferred to the ER by SYT1 and SYT3 at ER-PM CS in order to avoid membrane damage. This removal of DAG from the PM will be further enhanced by the stress-induced expansion of ER-PM CS marked by SYT1 and SYT3.

## ACKNOWLEDGMENTS

This work was supported by the Ministerio de EconomÍa y Competitividad, co-financed by the European Regional Development Fund (grant BIO2017-82609-R to M.A.B), the Marie Skłodowska-Curie actions (grant H2020-655366-IIF-PLICO to M.A.B and N.R-L), the Ministerio de Ciencia, Innovación y Universidades (grant PGC2018-098789-B-I00 to N.R-L), UMA-FEDER (grant UMA18-FEDERJA-154 to N.R-L) and by the Ministerio de EconomÍa, Industria y Competitividad (grant RYC-2013-12699 to N.R-L). J.P-S and S.G-H were funded by the Ministerio de EconomÍa y Competitividad in Formación del Personal Investigador Fellowship (BES-2012-052324) and (PRE2018-085284), respectively. R.P.H and J.A.N received support from the Biotechnology and Biological Sciences Research Council (BBSRC, UK) in the form of an Institute Strategic Programme Grant (BBS/E/C/000I0420). J.L. is supported by the Program of Introducing Talents of Discipline to Universities (111 Project, B13007). A.P.M and J.P-S were supported by the Shanghai Center for Plant Stress Biology (Chinese Academy of Sciences), Chinese 1000 Talents Program. A.R. was supported by the Natural Sciences and Engineering Research Council of Canada (NSERC-Discovery Grant RGPIN-2019-05568). Support was also provided by AEI/FEDER, UE (Grants BIO2016-79187-R and PID2019-106987RB-I00 to J.P-S) and by the European Research Council under the European Union’s Seventh Framework Programme (FP7/2007-2013) / ERC grant agreement n° 742985 to J.F. and T-Rex project number 682436 to D.V.D.).

We would also like to thank Lothar Willmitzer for the lipidomic analysis at the Max Planck Institute of Molecular Plant Physiology (Potsdam, Germany). We thank Manuela Vega from SCI for technical assistance in image analysis. We thank John R. Pearson and the Bionand Nanoimaging Unit, F. David Navas Fernández and the SCAI Imaging Facility and The Plant Cell Biology facility at the Shanghai Center for Plant Stress Biology for assistance with confocal microscopyThe FaFAH1 clone was a gift from Iraida Amaya Saavedra (IFAPA-Centro de Churriana, Málaga, Spain). The MAP-mTU2-SAC1 construct was provided by Yvon Jaillais (Laboratoire Reproduction et Développement des Plantes, Univ Lyon, France). The pGWB5 from the pGWB vector series, was provided by Tsuyoshi Nakagawa (Department of Molecular and Functional Genomics, Shimane University)

## AUTHORS CONTRIBUTION

M.A.B., N.R-L., and J.P-S. conceived the project. All authors contributed to the experimental design. N.R-L., J.P-S., A.E-dV., R.P.H., S.V., R.C., C.P-R., S.G-H., J.V., and V.A-S. performed the experiments and analyzed the data. M.A.B., N.R-L. and J.P-S. wrote the article. All authors reviewed and commented on the article.

## METHODS

### Plant Material

All Arabidopsis (*Arabidopsis thaliana*) plants in this study are Col-0 ecotype. Arabidopsis mutant lines used in this study are: *syt1* (AT2G20990) SAIL_775_A08 that has been described previously (Perez Sancho et al., 2015), *syt3* (AT5G04220) SALK_037933 (obtained from the Arabidopsis Biological Resource Center, Ohio State University) and *syt1/3* double mutant obtained by crossing *syt1* with *syt3*. The transgenic lines SYT1:SYT1-GFP (Perez Sancho et al., 2015), *syt1*/SYT1:SYT1-GFP (Lee et al, 2019) and 35S:GFP (Wang et al., 2019) were previously describe. Line syt1/SYT1:SYT1-GFP/1xPH^FAPP1^ was generated by crossing *syt1*:SYT*1-SYT1-GFP* with the PI(4)P marker P1xPH^FAPP1^ (Simon et al., 2016). Generation of transgenic lines *SYT1:MAPPER, SYT3:SYT3-GFP, 35S:SYT3-GFP* and *SYT3:GUS* is described in the section “Generation of Transgenic Plants.” Line *syt1*/35S:SYT3-GFP was obtained by crossing *syt1* with the *35S:SYT3-GFP* line.

### Plant Manipulation and Growth Conditions

Arabidopsis standard handling procedures and conditions were used to promote seeds germination and growth. First, Arabidopsis seeds were sterilized and cold treated for 3 days at 4°C. Next, seeds were sowed onto half-strength Murashige and Skoog (MS) agar solidified medium (0.6% [w/v] agar for horizontal growth and 1% [w/v] for vertical growth) containing 1.5% [w/v] Suc, unless otherwise stated. Plates were placed either vertically or horizontally in a culture chamber at 23 ± 1°C, under cool-white light (at 120 µmol photon m^−2^ s^−1^) with a long-day photoperiod (at 16-h light/8-h dark cycle) unless otherwise stated. When required, seedlings were transferred to soil after 7 days of *in vitro* growth and watered every 2 days. In soil, plants were grown in a mixture of organic substrate and vermiculite (4:1 [v/v]) under controlled conditions at 23 ± 1°C, 16-h light/ 8-h dark cycle (at 120 µmol photon m^−2^ s^−1^). Freshly harvested seeds were used for all phenotypic analyses.

### Plasmid Constructs

*SYT3:SYT3-GFP, 35S:SYT3-GFP* and *SYT3:GUS* constructs were generated using Gateway™. A 2210-bp SYT3 promoter fragment was cloned into pDONRP4P1R via a BP reaction (Invitrogen) to generate pEN-L4-proSYT3-R1. The genomic fragment of the SYT3 coding sequence without stop codon was amplified from Col-0 DNA using primers listed in Supplemental Table S7 and recombined into pDONR221 via a BP reaction (Invitrogen) to generate pEN-L1-SYT3g-L2. For the SYT3:SYT3-GFP and 35S:SYT3-GFP, we recombined respectively pEN-L4-proSYT3-R1 or pEN-L4-2-R1 (proCaMV35S) (Karimi et al., 2007) with pEN-L1-SYT3g-L2 and pEN-R2-F-L3,0 (GFP) (Karimi et al., 2007) into pKm43GW (Karimi et al., 2005). For SYT3:GUS construct, pEN-L4-proSYT3-R1 was recombined with pEN-L1-S-L2 (GUS) into pKm42GW (Karimi et al., 2005). The SYT1:MAPPER construct was generated by recombination of the previously described constructs pEN-L4-proSYT1-R1 and pEN-L1-MAPPER-L2 (Perez Sancho et al., 2015) with pEN-R2-empty-L3 and the destination vector pKm42GW.

Multisite Gateway™ cloning was used in the preparation of constructs for transient expression in *N. benthamiana*. The coding DNA sequence (CDS) without the stop codon of *SYT1, SYT3* and *DGK2* (AT5G63770) were PCR amplified using the primers listed in Supplemental Table S7 and cloned into the pENTR/ZEO vector using BP cloning kit (Invitrogen). All the pENTR clones were verified by sequencing. The pENTR vector with the CDS of *FaFAH1* (Sánchez-Sevilla et al., 2014) was a gift from Iraida Amaya. These pENTR clones in combination with the appropriate destination vectors (pDEST) were used to create the final Gateway-expression constructs by LR-reaction (Invitrogen). The *pGWB5* was used to generate *35S:SYT1-GFP* and *35S:SYT3-GFP* constructs. The pENTR vectors, pEN-L4-pUBQ10-R1 and pEN L2-mCherry-R3 were used with the pDEST vector pH7m34GW and the generated pEN-L1-L2 vectors to produce *UBQ10:SYT1-mCherry, UBQ10:SYT3-mCherry, UBQ10:DGK2-mCherry* and *UBQ10:FaFAH1-mCherry* constructs. The *SYT1:MAPPER* used for *N. benthamiana* experiments was the same as in the generation of *A. thaliana* lines. The construct MAP-mTU2-SAC1 is previously described (Simon et al., 2016).

### Generation of Arabidopsis Transgenic Lines

Expression constructs were transformed into *Agrobacterium tumefaciens* strain GVG3101::pMP90 through electroporation and confirmed by diagnostic PCR. The *SYT1:MAPPER, SYT3:SYT3-GFP, 35S:SYT3-GFP*, abd *SYT3:GUS* constructs were individually transformed into Arabidopsis Col-0 plants by floral dip to generate stable transgenic plants. SYT1:SYT1-GFP construct was transformed into the appropriate genotypes and then crossed to the single *syt1* mutant. T3 or T4 homozygous transgenic plants were used in this study. Line 35S:SYT3-GFP was crossed with line *syt1* to generate *syt1/*35S:SYT3-GFP.

### Transient Expression in *N. benthamiana*

Transient expression in *Nicotiana benthamiana* was performed using *Agrobacterium* strain (GV3101::pMP90) carrying the different constructs, together with the Agrobacterium expressing p19. Four to five-week-old *N. benthamiana* leaves were infiltrated with *Agrobacterium* into at the abaxial side of the leaf lamina. After infiltration, all plants were kept in the growth chamber and analyzed 2 days later. Agrobacteria cultures were grown overnight in Luria-Bertani medium containing rifampicin (50 mg/mL), gentamycin (25 mg/mL), and the construct-specific antibiotic. Cells were then harvested by centrifugation and pellets were resuspended in agroinfiltration solution (10 mM MES, pH 5.6, 10 mM MgCl_2_, and 1 mM acetosyringone), and incubated for 2 hr in dark conditions at room temperature. Agrobacterium strains were infiltrated at OD600 of 0.70 for the constructs and 0.25 for the p19 strain. For double infiltration experiments, Agrobacterium strains were infiltrated at OD600 of 0.40 for the constructs and 0.15 for the p19 strain.

### Arabidopsis eFP Browser Data Analysis

Gene expression level data from abiotic stress responses were retrieved from the Arabidopsis eFP Browser (Abiotic Stress Series) website (http://bar.utoronto.ca/efp/cgi-bin/efpWeb.cgi). Data used for the analysis were obtained from 18-day-old WT seedlings. Differential expression was calculated by dividing the expression value of each gene in a given abiotic stress by the corresponding control (fold change of abiotic stress relative to the mock). The abiotic stress gene expression response was calculated and the heatmap was created using Excel (Microsoft). In the heatmap, red represents induction and blue represents repression as response to the indicated hormone.

### Total RNA Extraction for RNA-seq

Three-week-old Arabidopsis rosettes grown in soil were used for total RNA extraction. Plant were treated for 24 h at 20°C (control) and 24 h at 4°C (cold) and rosette tissue was grounded to a fine powder in liquid nitrogen. Approximately 100 mg of ground tissue per sample was extracted using the Kit Pure link RNA Mini Kit (Ambion) following the manufacturer’s instructions. RNA samples (8 mg per sample) were treated with 1µL of Turbo DNAse (Ambion) at 37°C for 35 minutes. One mg of DNAse treated RNA samples were purified using the RNA Clean and Concentrator-25 (Zymo Research) according to the manufacturer’s instructions. Samples were measured in a Bioanalyzer to check the RIN.

### Total RNA Extraction and Quantitative PCR Analysis

Seven-day-old whole seedlings (20 seedlings per biological replicate) were used for total RNA extraction of 6 h and 24 h cold treatment experiments. Three-week-old plants were used for total RNA extraction of adult plants cold-acclimated during 1, 3 or 7 days. Plant tissue was grounded to a fine powder in liquid nitrogen. Approximately 100 mg of grounded tissue per sample was homogenized in 1 mL of the commercial reagent TRIsure (Bioline), and total RNA was extracted following the manufacturer’s instructions. RNA samples (10 mg per sample) were treated with TurboDNA-free DNase (Ambion), and 1mg of RNA per sample was run on a 1% agarose gel to confirm RNA integrity. First-strand cDNA was synthesized from 1 mg of RNA using the iScript cDNA synthesis kit (Bio-Rad), according to the manufacturer’s instructions. cDNAs were amplified in triplicate by quantitative PCR (qPCR) using SsoFast EvaGreen supermix (Bio-Rad) and the Rotor-Gene Q cycler (Qiagen). The amplification protocol includes an initial step at 95°C for 2 min, followed by 40 cycles of 5 s at 95°C, 15 s at 58°C and 20 s at 72°C. The relative expression values were determined using ACTINE2 as a reference. Primers used for qPCR are listed in Supplemental Table S7.

### RNA-sequencing

Transcriptome analyses were performed at the Genomics Core Facility, Shanghai Center for Plant Stress Biology, CAS. Four biological replicates of each genotype were used. Total RNA (1 μg) from each sample was used for library preparation with NEBNext® Poly(A) mRNA Magnetic Isolation Module (New England BioLabs) and NEBNext Ultra II Directional RNA Library Prep Kit for Illumina (New England BioLabs) following the manufacturer’s instructions. Prepared libraries were assessed for quality using NGS High-Sensitivity kit on a Fragment Analyzer (AATI) and for quantity using Qubit 2.0 fluorometer (Thermo Fisher Scientific). All libraries were sequenced in paired-end 150 bases protocol (PE150) on an Illumina HiSeq sequencer.

### Meta-analysis (GO-Terms)

Functional enrichment analyses of the Biological Process ontology were performed using VirtualPlant (Katari et al., 2010).

### Extraction of total proteins from Arabidopsis

Arabidopsis tissue was grounded to a fine powder in liquid nitrogen. Approximately 100 mg of grounded tissue per sample was used for total protein extraction. Denatured protein extracts were obtained by homogenizing and incubating plant material in 2⨯ (2 uL of buffer per mg of tissue) Laemmli buffer (125 mM Tris-HCl, pH 6.8, 4% [w/v] SDS, 20% [v/v] glycerol, 2% [v/v] b-mercaptoethanol, and 0.01% [w/v] bromophenol blue) for 20-30 min at 70°C (vortexing during the incubation). Extracts were centrifuged for 5 min at 20,000 g at 10-15°C and supernatants were recovered. The total protein extracts from supernatant were separated in a 10% SDS-PAGE gel and analyzed as described in the section “Immunoblot.”

### Immunoblot

Proteins separated by SDS-PAGE polyacrylamide gel electrophoresis were electroblotted using Trans-blot Turbo Transfer System (Bio-Rad) onto polyvinylidene difluoride (PVDF) membranes (Immobilon-P, Millipore) following instructions by the manufacturer (preprogramed protocols optimized for the molecular weight of the proteins of interest). PVDF membranes, containing electroblotted proteins, were then incubated with the appropriate primary antibody followed by the appropriate secondary peroxidase-conjugated antibody. The following primary antibodies were used for detection of epitope-tagged proteins: mouse monoclonal anti-GFP clone B-2 (1:1000, catalog no. sc-9996, Santa Cruz Bio-technology) and rabbit polyclonal anti-SYT1 antibody (1:5000). The secondary antibodies used in this study were as follows: anti-mouse IgG whole molecule-Peroxidase (1:80,000; catalog no. A9044, Sigma-Aldrich) and anti-rabbit IgG whole molecule-Peroxidase (1:80,000; catalog no. A0545, Sigma-Aldrich). Proteins and epitope-tagged proteins on immunoblots were detected using the Clarity ECL Western Blotting Substrate or SuperSignal West Femto Maximum Sensitivity Substrate according to the manufacturer’s instructions, and images of different time exposures were acquired using the Chemidoc XRS1System (Bio-Rad). SDS-PAGE polyacrylamide gels and immunoblotted PVDF membranes were stained with Coomassie Brilliant Blue R 250 to confirm equal loading of the different samples in a given experiment.

### Co-Immunoprecipitation in Arabidopsis

For Co-IP experiments, 13-day-old Arabidopsis plants were grounded to fine powder in liquid nitrogen. Approximated 0.5 g of grounded leaves per sample were used and total proteins were then extracted with extraction buffer [50 mM Tris-HCl, pH 7.5; 150 mM NaCl; 10% glycerol; 10 mM EDTA, pH 8; 1 mM NaF; 1 mM Na2MoO4·2H2O; 10 mM DTT; 0.5 mM PMSF; 1% (v/v) P9599 protease inhibitor cocktail (Sigma); Nonidet P-40, CAS: 9036-19-5 (USB Amersham life science) 0.5% (v/v)] added at 2 mL/g of powder using an end-over-end rocker for 30 minutes at 4°C. Samples were centrifuged 20 minutes at 4°C at 9,000g. Supernatants were filtered by gravity through Poly-Prep Chromatography Columns (#731-1550 Bio-Rad) and 100 _μ_L was reserved for immunoblot analysis as input. The remaining supernatants were diluted in (1:1 dilution) with extraction buffer without Nonidet P-40, so the final concentration of detergent (Nonidet P-40) was adjusted to 0.25% (v/v) to avoid unspecific binding to the matrix as recommended by the manufacturer. Protein samples were then incubated for 2 hours at 4°C with 15 _μ_L GFP-Trap coupled to agarose beads (Chromotek) in an end-over-end rocker. Following incubation, the beads were collected and washed four times with the wash buffer (extraction buffer without detergent). Finally, beads were resuspended in 75 _μ_L of 2x concentrated Laemmli Sample Buffer and heated at 70°C for 30 minutes to dissociate immunocomplexes from the beads. Total (input), immunoprecipitated (IP) and Co-Immunoprecipitated (CoIP) proteins were separated in a 10% SDS-PAGE gel and analyzed by Immunoblot.

### PAO treatment

Six-day-old seedlings of SYT1-GFP/1xPH^FAPP1^ were incubated in wells containing 60 µM of PAO (Sigma, PAO stock solution 60 mM in DMSO) in one half strength liquid MS for 4 hours. The mock condition corresponds to the incubation of plants in wells supplemented with the same volume of DMSO and treated for the same time. Roots were then subjected to imaging using Confocal Laser Scanning Microscopy.

### Cold and salt treatments for cell viability in roots

Previous to cold treatment, 24-well plates loaded with liquid one tenth strength MS solution was placed on ice for 30 minutes. Six-day-old seedlings of the phenotypes in study (WT, *syt1, syt3, syt1/3, syt1/*SYT1:SYT1-GFP and *syt1*/35S:SYT3-GFP) were incubated in the wells containing the mentioned liquid media at 8°C at different times between 0 and 30 min. Seedlings were then stained in 1/10 MS containing 10 µg/ml of FDA (Sigma, PDA stock solution 5 mg/ml in DMSO) for 5 min. Seedling were then washed once and imaging was performed. In all cases, roots were imaged within a 5-minute time frame window around the indicated time. For salt treatment, 4-day-old seedlings of WT, *syt1, syt3* and *syt1/3* were incubated in 150 mM KCl in one tenth MS strength for 1 hour. Then, they were stained as explained above and imaging was performed. In both cases, at least 10 plants of each phenotype were used (and the experiment was repeated twice).

### Cold treatments for SYT1-GFP and SYT3-GFP dynamics

Arabidopsis seedlings expressing *SYT1:SYT1-GFP* or *35S:SYT3-GFP* were grown vertically in half-strength MS agar solidified medium under long-day photoperiod and 23°C for 7 days. For lon exposure cold treatment, plates were transferred to a temperature programmable chamber and kept at 4°C for the specified times. Control plates kept growing in control conditions. For short-time cold exposure, seedlings were covered with liquid half-strength MS pre-chilled at 4°C and the plates were transferred to a box with ice for the specified times. Mock seedlings were covered with liquid half-strength MS at 23°C for the specified times while control seedlings were undisturbed. Mounting of the cold-treated seedlings on the microscope slides was done under cold conditions and with pre-chilled water. In all cases, roots were imaged between a 3-4-minute time frame window after taken out from the MS plate.

### Immunostaining of Arabidopsis roots

Five-day-old root tips were used, and all working solutions were prepared in microtubule stabilization buffer (50 mM PIPES, 5 mM EGTA, and 5 mM MgSO_4_ [pH 7.0]). Sample preparation followed the basic protocol described in Sauer and Friml (2010). For immunodetection, a polyclonal rabbit anti-SYT1 anti-body (1:1,000) was incubated overnight at 4°C plus two additional hours at 37°C. The root tips were incubated then for 1 h with TRITC-conjugated AffiniPure Donkey anti-Rabbit IgG (1:200; Jackson Immunoresearch), and mounted on microscopy slides using the glycerol-based AF1 Mountant Solution (Citifluor).

### Electrolyte Leakage Measurements

One-week-old seedlings grown in half-strength MS plates were transferred to tubes containing 5 ml of 20% PEG and incubated at 25°C for the indicated time points. Afterwards, seedlings were carefully removed, washed with deionized water, and placed in new tubes containing 5 mL of deionized water. The tubes were shaken at 120 rpm for 3 hours at 25°C and the conductivity of the solutions was measured. The tubes were then autoclaved and after cooling down to room temperature, the conductivities of the solutions were measured again. The percentage of electrolyte leakage was calculated as the percentage of the conductivity before autoclaving over that after autoclaving. Three independent experiments were performed.

### Generation and Purification of GST Fusion Proteins

RT-PCR of WT mRNA was used to produce GST fusion clones in pGEX-KG corresponding to the following SYT3 amino acid sequences: 244 to 402, corresponding to the SYT3-C2A domain, and 403 to 540, corresponding to the SYT3-C2B domain. Briefly, proteins expressed in BL21-RP were released by sonication and incubated with glutathione agarose beads (0.3 mL/L culture) overnight at 48°C. Proteins were washed on the agarose beads four times with 10 mL PBS, 1 mM EDTA, and 0.1 g/L PMSF. Purified protein was eluted from the agarose with 2.5 mL 100 mM Tris, pH 8.0, and 40 mM glutathione and concentrated to 0.2 mL by centricon centrifugation (Millipore). Protein concentrations were determined by the method of Bradford with protein dye reagent concentrate from Bio-Rad, using BSA as standard.

### Phospholipid Binding Assays

Phospholipid binding to isolated C2A and C2B domains of SYT3 was assessed by a centrifugation assay as described previously (Schapire et al., 2008). Briefly, phospholipids (PS/PC = 25/75, w/w) (Sigma-Aldrich) in chloroform were dried as a thin layer under a stream of nitrogen. Dried lipids were resuspended in buffer A (100 mM NaCl; 50 mM HEPES, pH 6.8; 4 mM EGTA) by vortexing for 20 min. Large multilamellar vesicles were collected by centrifugation for 20 min at 20,800*g* and resuspended in buffer A with various concentrations of free Ca^2+^ and used within 1 h. Calcium concentrations were calculated using the WEBMAXC STANDARD, RRID:SCR_003165). Purified soluble recombinant GST-C2 domains (6 _μ_<u>g</u>) and vesicles (100 _μ_g of phospholipids; total volume = 1 mL) were incubated for 15 min at 27°C with shaking at 250 rev/min on a platform shaker. Large multilamellar vesicles and bound protein were isolated by centrifugation for 10 min at 20,800*g* at 4°C. Pellets were washed three times with 0.5 mL of the corresponding incubation buffer, and the bound protein was analyzed by SDS-PAGE and densitometry using a Bio-Rad GS670 scanning densitometer of the Coomassie Brilliant Blue– stained gel.

### Modeling of SYT3 C2 domains

Analysis of torsional angles using Rampage (Lovell et al., 2003) and Ramachandran plots were generated (Figure S2). As a result, very robust C2A^244-402^ and C2B^403-540^ 3D models with a high confidence match (>99%) were generated. Three Ca^+2^ coordination pockets with a high confidence score (Cs>0.5) were identified for C2A2 while no Ca^+2^ coordination pockets were identified for C2B.

### Total lipids extraction and analysis by Liquid Chromatography-Mass Spectrometry

Lipid extraction was performed as previously described (Giavalisco et al., 2011)) at Max Planck Institute of Molecular Plant Physiology. In brief, 100 mg of each biological replicate was extracted in 1 mL of a pre-cooled mixture of methanol/methyl-tert-butyl-ether/water (1:3:1) and shaking in a cooled sonic bath for 10 min. Then, 500 µl of water:methanol (3:1) was added. This led to the formation of two phases. A fixed volume of lipid phases was transferred to a Eppendorf tube and vacuum-dried to dryness. Chromatographic separation was performed using a Waters Acquity UPLC system connected to a Exactive Orbitrap (Thermo Fischer Scientific) via a heated electrospray source (Thermo Fischer Scientific). Processing of chromatograms, peak detection and integrations were done with REFINER MS 7.5 (GeneData; http://www.genedata.com). The obtained features (*m/z* at a certain retention time) were queried against an in-house lipid database for further annotation.

### Plasma membrane isolation

Plasma membrane purification was carried out as described previously (Bernfur et al., 2013) with some modifications. All steps in the preparation procedure were performed at 4°C or on ice. Fifteen grams of leaves were homogenized with a knife blender in 40 ml homogenization buffer: 0.33 M sucrose, 50 mM MOPS-KOH, pH 7.5, 5 mM EDTA, 5mM EGTA, 20mM NaF and including 0.6% (w/v) polyvinylpoly-pyrrolidone, 5 mM ascorbate, 4mM salicylhydroxamic acid (SHAM), 150µM 4-(2-aminoethyl)-benzene-sulfonyl fluoride (AEBSF) and 5 mM dithiothreitol (DTT) that were added immediately before use. The homogenate was filtered through a nylon mesh (200 µm) and phenylmethanesulphonyl-fluoride (PMSF) was added to a final concentration of 1 mM. The filtrate was centrifuged at 10,000 g for 15 min, the pellet was discarded, and the supernatant was centrifuged at 100,000 g for 2 h at 4°C. The microsomal pellet was resuspended in 1.5-2.0 ml resuspension buffer: 0.33 M sucrose, 5 mM K-phosphate, pH 7.8, 0.1 mM EDTA, and 1 mM DTT freshly added Final weight of the resuspended pellet should be 3.0 g. To produce an aqueous polymer two-phase system with a final weight of 12 g, the resuspended pellet (3 g) was added to a 9.00 g phase mixture producing a 6.1% (w/w) dextran 500, 6.1% (w/w) polyethylene glycol 3350, 0.33 M sucrose, 5 mM K-phosphate, pH 7.8, 3 mM KCl phase system. Further purification of the plasma membranes using the aqueous polymer two-phase system was performed according to (Larsson et al., 1994). The final upper phase, highly enriched in plasma membranes, was diluted two-fold with 0.33 M sucrose, 5 mM K-phosphate, pH 7.8, 0.1 mM EDTA before centrifugation at 100,000 g for 2 h. The plasma membrane pellet was resuspended in 100 µl resuspension buffer plus 5 mM KCl and stored at - 80 ° C until use.

### Protein large-scale immunoprecipitation for lipidomics

Arabidopsis lines *syt1/SYT1:SYT1-GFP* and *SYT1:MAPPER* were used for this analysis. Four-week-old Arabidopsis rosettes were ground to fine powder in liquid nitrogen. Fifteen grams of ground leaves per sample were used and total proteins were then extracted with extraction buffer [50mM Tris-HCl pH 7.5, 150 mM NaCl, 10 % glycerol; 16mM CHAPS, 10 mM EDTA pH 8.0, 1 mM Na2MoO4·2H2O, 1 mM NaF, 10 mM DTT, 0.5 mM PMSF; and 1% (v/v) P9599 protease inhibitor cocktail (Sigma)] added at 2 mL/g of powder and mixed by pipetting and with an end-over-end rocker for 30 minutes at 4°C. Samples were centrifuged at 9000 g for 20 min at 4°C. Supernatants were filtered by gravity through Poly-Prep Chromatography Columns (#731-1550 Bio-Rad) and 100 _μ_L was reserved for immunoblot analysis as input. The remaining supernatants were diluted (1:1 dilution) with extraction buffer without CHAPS. Protein samples were then incubated for 2 hours at 4°C with 150 _μ_L GFP-Trap coupled to agarose beads (Chromotek) in an end-over-end rocker. Following incubation, beads were collected and washed three times with wash buffer (extraction buffer without detergent) and one more time with wash buffer with 500 mM NaCl. An aliquot of the beads (approx. 1/10) was saved for co-inmunoprecipitation analysis. For these samples, proteins were stripped from these beads by boiling in SDS loading buffer for 10 mins. Immunoprecipitated proteins were separated on SDS-PAGE acrylamide gels and detected by immunoblot. Additionally, lipids of immunoprecipitated proteins in the rest of the beads were extracted and analysed. Lipids were extracted as previously described using the method of Bligh and Dyer (see below).

### Plasma membrane, SYT1-GFP and MAPPER samples lipid extraction and analysis

Lipids were extracted as previously described using the method of Bligh and Dyer. Briefly, extraction of lipids from immunoprecipitated proteins in GFP-beads or plasma membrane samples was performed by the addition of 3.75 volumes of chloroform:methanol (1:2 v/v), followed by sonication for 1-2 min and vortex during 15-20 min. Then 1.25 volumes of chloroform were added, samples were vortexed for 1 min and 1.25 volumes of 0.5 % acetic acid in 500 mM of chloroform were added, followed by 1 min of vortexing. The sample was centrifuged at low speed (300 x g) using a tabletop centrifuge for 5 min; then the bottom layer was gently recovered. Two upper phase re-extractions were done by adding 1.875 vol of chloroform, followed by vortex and centrifugation as above. Lipids were dried by adding gas N_2_. Quantitative analyses of lipids (at Rothamsted Research, UK), including neutral (diacylglycerols, DAG) and polar lipids (phosphatidylcholine (PC), phosphatidylethanolamine (PE), phosphatidylinositol (PI), phosphatidylglycerol (PG), Lysophosphatidylcholine (LPC), monogalactosyldiacylglycerol (MGDG) and digalactosyldiacylglycerol (DGDG)) lipids were carried out using electrospray ionization tandem triple quadrupole mass spectrometry (API 4000 QTRAP; SCIEX; ESI-MS/MS) as described previously (Guo et al., 2019). The internal standards were supplied by Avanti (Alabama, USA), incorporated as 8 pmol 13:0-LPC, 0.086 nmol di24:1-PC, 0.080 nmol di14:0-PE, 0.05 nmol di18:0-PI, 0.080 di14:0-PG, 0.03 nmol di18:0-PS and 0.03 nmol di14:0-PA. The standards dissolved in chloroform and 25 _μ_L of the samples were combined with chloroform/methanol/300 mM ammonium acetate (300:665:3.5 v/v/v) to make a final volume of 1 ml. The lipid extracts were infused at 15 _μ_l/min with an autosampler (HTS-xt PAL, CTC-PAL Analytics AG). Data acquisition and acyl group identification of the polar lipids were carried out as described in (Ruiz-Lopez et al., 2014) with modifications. For quantifying DAGs, 25 _μ_l of lipid extract and 0.043 nmol 18:0-20:4-DAG (Nu-Chek-Prep) were combined with chloroform/methanol/300 mM ammonium acetate (24:24:1.75 v/v/v), to final volumes of 1 ml for direct infusion into the mass spectrometer. DAGs were detected as [M + NH4]+ ions by a series of different neutral loss scans, targeting losses of FAs. Full documentation of lipid profiling data is provided in Supplementary Table S5 and S6.

### Confocal Imaging of Arabidopsis and *N. benthamiana*

Living cell images were obtained using two different inverted Leica TCS SP5 II confocal laser-scanning microscope, and a Nikon Eclipse Ti based Andor Revolution WD spinning-disk confocal microscope. One of the Leica TCS SP5 II laser-scanning microscopes was equipped with GaAsp HyD detectors. Both were equipped with HCX PL Apo 40x/ NA1.3 and HCX PL Apo 63x/ Na1.4 oil immersion objective lenses. The software Leica LAS X was used for image acquisition. The Andor Revolution WD microscope was equipped with a CSU-W1 spinning disc head (Yokogawa, Tokyo, Japan), an Andor iXon Ultra 888 (EMCCD) camera and a Nikon Apo TIRF 60x/NA 1.49 oil immersion objective lens. The software Leica LAS X was used for image acquisition. Andor iQ3.6. All microscopes were equipped with a 488 nm laser for GFP/Fluorescein excitation, and the Leica SP5 II were also equipped with a 561 nm He-Ne laser for mCherry/ TRITC excitation.

For imaging on *N. benthamiana* leaves, 2 days after infiltration leaf-discs were excised from the leaves immediately before visualization under the Leica TCS SP5 II microscope with 40x lens and the lower epidermis of the leaf was 3D imaged from the equatorial plane until the cell surface with 600 nm spacing. GaAsp HyD detectors were used to improve the signal detection. For colocalization, sequential line scanning mode was used to separate signals. Cortical plane images are a maximum Z-projection of several planes from the cell surface until a plane where cells are close but still not touching the neighbors (to ease identification of individual cells). Equatorial plane images are single plane images.

Imaging of SYTs patterns in Arabidopsis in control and cold conditions -long treatments- and immunostained Arabidopsis roots, was done with the Leica TCS SP5 II microscope. Imaging of SYTs patterns in Arabidopsis in mock and cold conditions -short treatments-, was done with the Andor Revolution WD spinning-disk confocal microscope. Cotyledons from 7-days-old seedlings were excised immediately before visualization and the epidermal cells in the adaxial side were imaged as indicated for *N. benthamiana*. Immunostained roots in Arabidopsis were visualized under the 63x lens and sequential line scanning mode was used.

Imaging of FDA stained roots was performed with the Leica TCS SP5 II microscope, using the 40x lens. Z-stacks were taken starting from the surface of the roots and ending 25-30 µm deep inside the root (1 µm spacing).

For the PAO treatment, single plane images were taken with the Leica TCS SP5 II microscope using the 40x objective. Sequential line scanning mode was used to capture the mCherry and GFP signals to avoid crosstalk.

### Image analysis

All image analyses were performed using FIJI (Schindelin et al., 2012). For the analysis of FDA stained roots, Z-projections (Max intensity) were obtained from FDA stained confocal stacks of Arabidopsis roots. All images were selected to the same size using the root tip as reference. Then they were thresholder and converted into binary images (using Auto Threshold, method Moments). In this way, the alive cells are shown in white, while the dead ones in black. Five different Regions of interests (ROIs, size 73,48 ⨯ 73,48 µm2) were chosen for each image and the area of the white signal was measured. As an example, a ROI where all cells are alive will have an area of 5400 µm2, while other in which all cells are dead will score 0. Data were normalized to the ROIs where all the cells were alive (it means, divided by 5400) and expressed as a percentage. For the analysis the images of short-time cold treated and mock plants,

images were first deconvolved using the Autoquant X3 Software (MediaCybernetics). For the quantification of contact sites, z-projections were obtained according to the following macro:

run(“Enhance Contrast…”, “saturated=0.005 normalize process_all”);

run(“Subtract Background…”, “rolling=20 stack”);

run(“Green”);

waitForUser (“make a z projection”);

run(“Subtract Background…”, “rolling=20 stack”);

run(“Smooth”);

waitForUser (“Do you like the result?”)

saveAs(“Tiff”);

close();

Three different Regions of interests (ROIs, size 150 x 150 px) were chosen for each image trying to avoid internal filamentous signals sometimes present. Intensity maximas (contact sites) were identified in each ROI using the following macro:

run(“Duplicate…”, ““);

run(“8-bit”);

run(“Subtract Background…”, “rolling=20”);

run(“Smooth”);

run(“Smooth”);

run(“Enhance Contrast…”, “saturated=0.1 normalize”);

run(“Find Maxima…”, “noise=20 output=Count exclude”);

### GUS staining assay

Whole five-day-old seedlings were immersed in histochemical GUS staining buffer (100 mM NaPO4, pH 7; 0.5 mM K3[Fe(CN)6; 0.5 mM K4[Fe(CN)6]; 20% methanol; 0.3% Triton X-100; and 2 mM 5-bromo-4-chloro-3-indoxyl-b-D-glucuronide cyclohexylammonium) in 12-well plates (5 seedlings per well), vacuum infiltrated (60 cm Hg) for 10 min, and then wrapped in aluminum foil and incubated at 37°C for 12 h. Samples were then washed with a mix of water and ethanol with increasing concentration of ethanol (25 %, 50 %, 75 %, 95 %) and finally several times with 95% ethanol until complete tissue clarification. Samples where then rehydrated by gradually reducing the ethanol concentration in the solution (95 %, 75 %, 50 %, 25 %). Samples were mounted in the microscopy slides in 50% glycerol and photographed using the Nikon AZ100 Multizoom microscope system.

