## Supplemental Figures for "Synaptotagmins Maintain Diacylglycerol Homeostasis at Endoplasmic Reticulum-Plasma Membrane Contact Sites during Abiotic Stress"

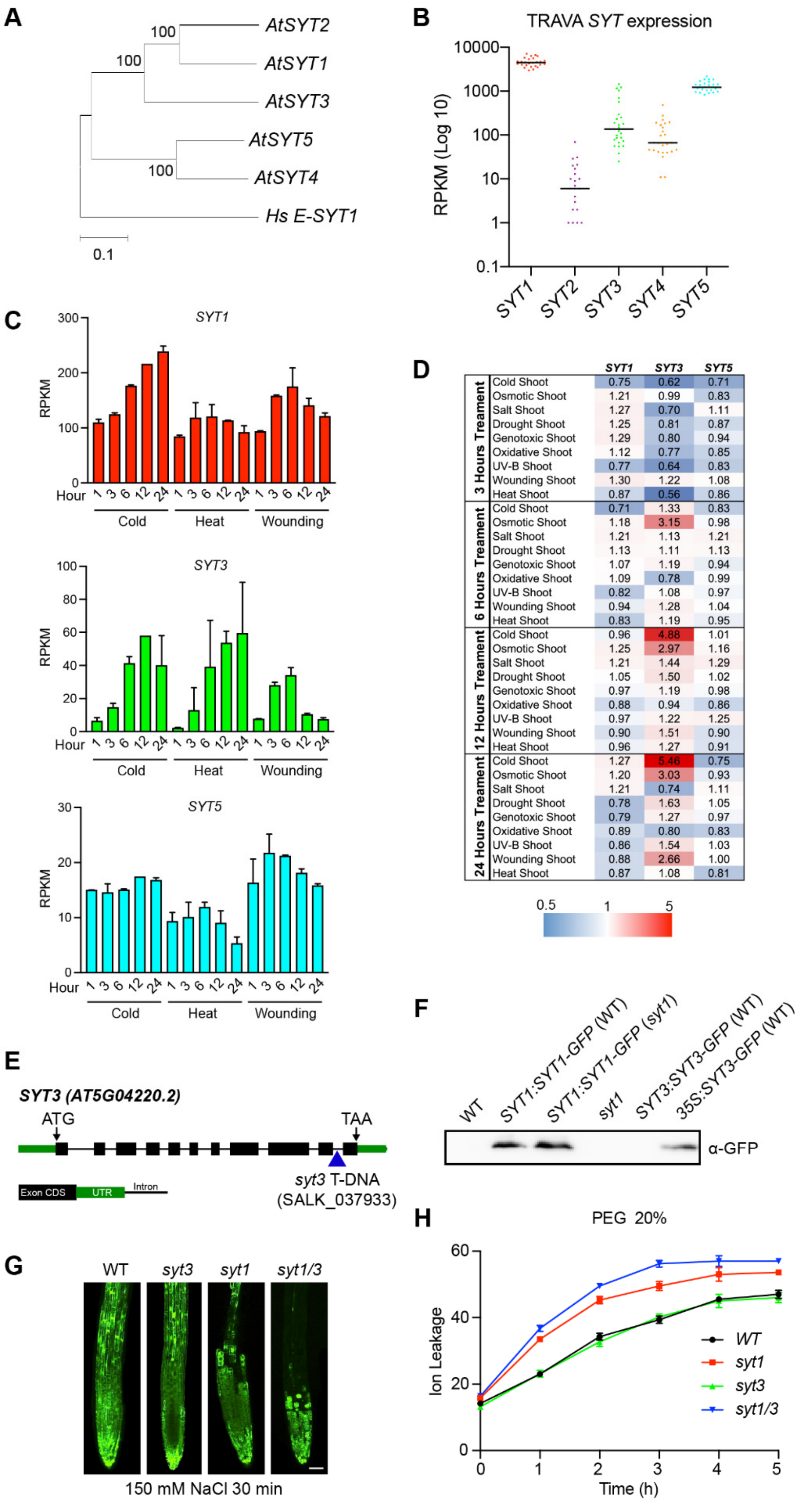


**Figure S1. *SYT1* and *SYT3* are induced under abiotic stress (related to Figure 1).**

**(A)** Phylogenetic clustering of the Arabidopsis protein coding sequence of SYT1, SYT2, SYT3, SYT4, SYT5 and the *Homo sapiens* E-Syt1. Phylogenetic tree was inferred using the Neighbor-Joining method. The optimal tree with the sum of branch length = 3.26 is shown. The tree is drawn to scale, with branch lengths in the same units as those of the evolutionary distances used to infer the phylogenetic tree. The evolutionary distances were computed using the p-distance method and are in the units of the number of amino acid differences per site. This analysis involved 11 amino acid sequences. All ambiguous positions were removed for each sequence pair (pairwise deletion option). There were a total of 1752 positions in the final dataset. Evolutionary analyses were conducted using MEGA X (Kumar et al., 2018).

**(B)** Gene expression profiles based on RNA-seq analysis of *SYT1*, *SYT2*, *SYT3*, *SYT4*, and *SYT5* in vegetative tissues at different developmental stages from TRAVA database http://travadb.org/). Each dot represent a RNAseq value and the bar represent the median.

**(C)** Gene expression profiles based on RNA-seq analysis of *SYT1*, *SYT3,* and *SYT5* in eFP-seq Browser after various abiotic stresses (https://bar.utoronto.ca/eFP-Seq_Browser/).https://bar.utoronto.ca/eFP-Seq_Browser/).

**(D)** Heatmap representing the expression responses to abiotic stress of *SYT1*, *SYT3* and *SYT5*. Expression levels genes are represented as the fold-change relative to the control. Red represents a gene that is induced, and blue represents a gene that is repressed in response to the indicated abiotic stress. Gene expression data was retrieved from the Arabidopsis eFP Browser (http://bar.utoronto.ca/efp/cgi-bin/efpWeb.cgi.

**(E)** T-DNA insertion site in *syt3-1* with exons shown as black boxes and intron as a line. The position of the start and stop codons are indicated.

**(F)** Protein accumulation in 10-day-old seedlings expressing *SYT1-GFP* (native promoter) or *SYT3-GFP* (native promoter or 35S promoter). Arabidopsis WT and *syt1* seedlings were used as control. Seedlings were grown under long-day photoperiod and 23°C for 10 days and whole seedlings were used for protein extraction. Proteins were detected by immunoblot with anti-GFP antibody. Experiment was repeated twice with similar results.

**(G)** Cell viability of root cells under NaCl stress determined by staining with fluorescein diacetate (FDA). *syt1* shows an increased sensitivity when compared to WT and *syt3* roots and *syt1/3* shows an increased sensitivity compared to *syt1*. Scale bar 50 μm.

**(H)** Ion leakage from 1-week-old seedlings treated with 20% PEG at different time points. Data are means of three independent assays. All mean values are significantly different between the WT and syt1 and between syt1/3 and syt1 ($p < 0.05$).

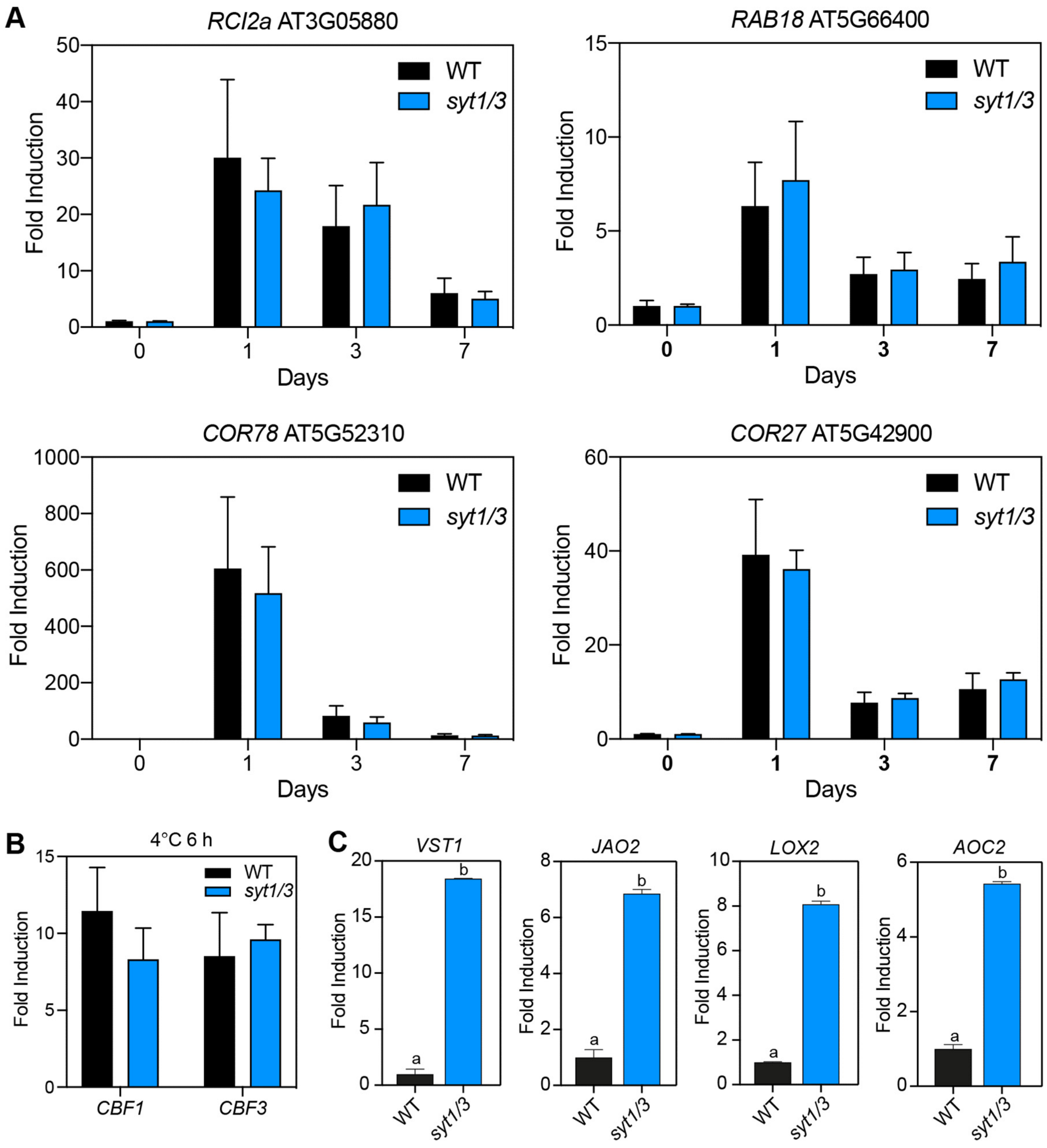


**Figure S2. Gene expression analyses in WT and *syt1/3* (related to Figure 2).**

**(A)** Cold acclimation marker genes *RCI12*, *RAB18*, *COR78* and *COR27* respond normally to cold in *syt1/3* plants. Arabidopsis WT and *syt1/3* plants were grown on soil under long-day photoperiod at 23°C and then transferred to 4°C for 1, 3 or 7 days. Relative expression level of the marker genes was measure by RT-qPCR. For each gene and each condition, the expression was normalized to the expression of *ACTIN2*. No significative differences between WT and *syt1/3* were found for any of the genes or time points analyzed.
**(B)** The *CBF1* and *CBF3* transcription factors show wilt type responses to cold in *syt1/3* seedlings. Arabidopsis WT and *syt1/3* seedlings were grown under long-day photoperiod and 23°C for 7 days and then plates were transferred to 4°C for 6 h or kept under control conditions. The relative expression level of the *CBF1* and *CBF3* genes was measured by RT-qPCR. For each gene, the expression with cold treatment was first normalized to the expression of *ACTIN2* and represented relative to its normalized expression in control conditions. No significative differences of *CBF1* and *CBF3* expression were found between WT and *syt1/3.*
**(C)** The *VST1*, *JAO1*, *LOX2*, *AOC2* expression is induced in *syt1/3* in response to cold. Three-week-old Arabidopsis WT and *syt1/3* plants were treated for 24 h at 4°C (cold) and rosette tissue was used for total RNA extraction and qPCR analysis. Lowercase letters indicate statistical differences between mutant vs WT determined by t-student statistical test ($p < 0.05$). Data represent mean values, error bars are SD, n=3.

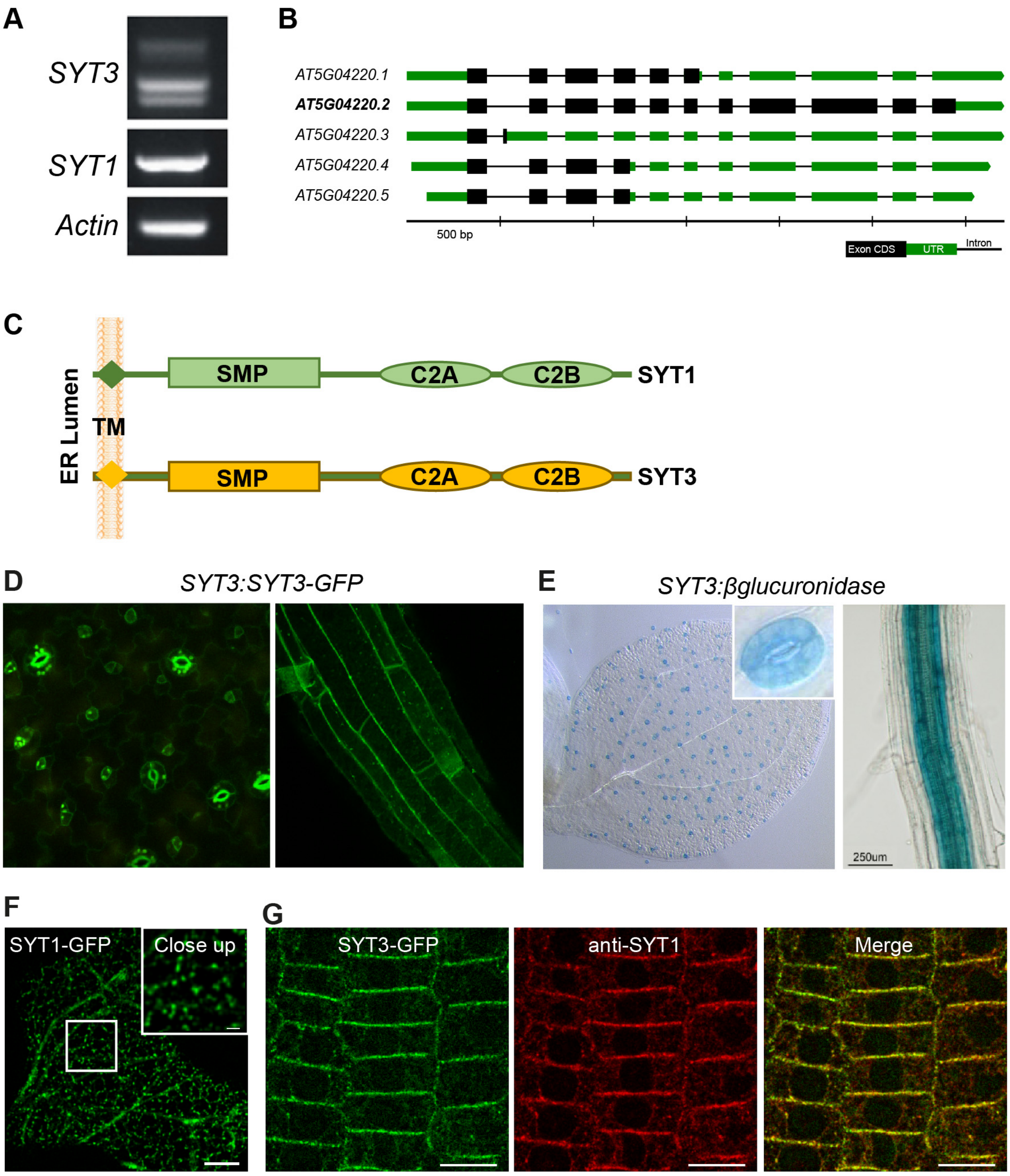


**Figure S3. *SYT3* alternative splicing, promotor expression and SYT3 subcellular localization (related to Figure 3).**

**(A)** RT-PCR analysis of WT plants showing the presence of alternatively spliced mRNAs variants with the smallest transcript encoding a full SYT3 protein.
**(B)** Gene model of *SYT3* alternative splice variants. Taller blocks (in black) show translated regions and thin blocks (in green) represent untranslated regions. The analyses of *SYT3* alternative splicing was performed using Integrated Genome Browser and is based in public available RNA-Seq data sets.
**(C)** Diagram of the structure of *Arabidopsis* SYT1 (top) and SYT3 (bottom). SYT1 and SYT3 are predicted to have a transmembrane domain, a SMP domain, and two C2 domains.
**(D)** Confocal images showing the localization of SYT3-GFP in the root (right panel) and in epidermal cells of the cotyledon (left panel) of 6-day-old Arabidopsis seedlings expressing *SYT3:SYT3-GFP* grown vertically in half-strength MS agar solidified medium under long-day photoperiod and 23°C.
**(E)** Images after histochemical GUS staining of cotyledon (left panel) and root (right panel) of 6-day-old Arabidopsis seedlings expressing *SYT3:GUS* grown vertically in half-strength MS agar solidified medium under long-day photoperiod and 23°C.

**(F)** Confocal image showing the localization of SYT1-GFP in epidermal cells of the cotyledon of 6-day-old Arabidopsis seedlings expressing *SYT1:SYT1-GFP* grown vertically in half-strength MS agar solidified medium under long-day photoperiod and 23°C. The boxed region is magnified in the inset (close up). Scale bar 10 μm and 2 μm for the close-up inset.
**(G)** SYT3-GFP colocalizes with endogenous SYT1 in Arabidopsis roots. Five-day-old Arabidopsis seedlings expressing *35S:SYT3-GFP* grown under long-day photoperiod and 23°C were fixed with 4 % PFA and permeabilized. The endogenous SYT1 protein was immunodetected in the root tip with a rabbit anti-SYT1 primary antibody and an anti-rabbit TRITC conjugated secondary antibody. Images of the individual channels as well as the merged image are shown.
Scale bar 10 μm.

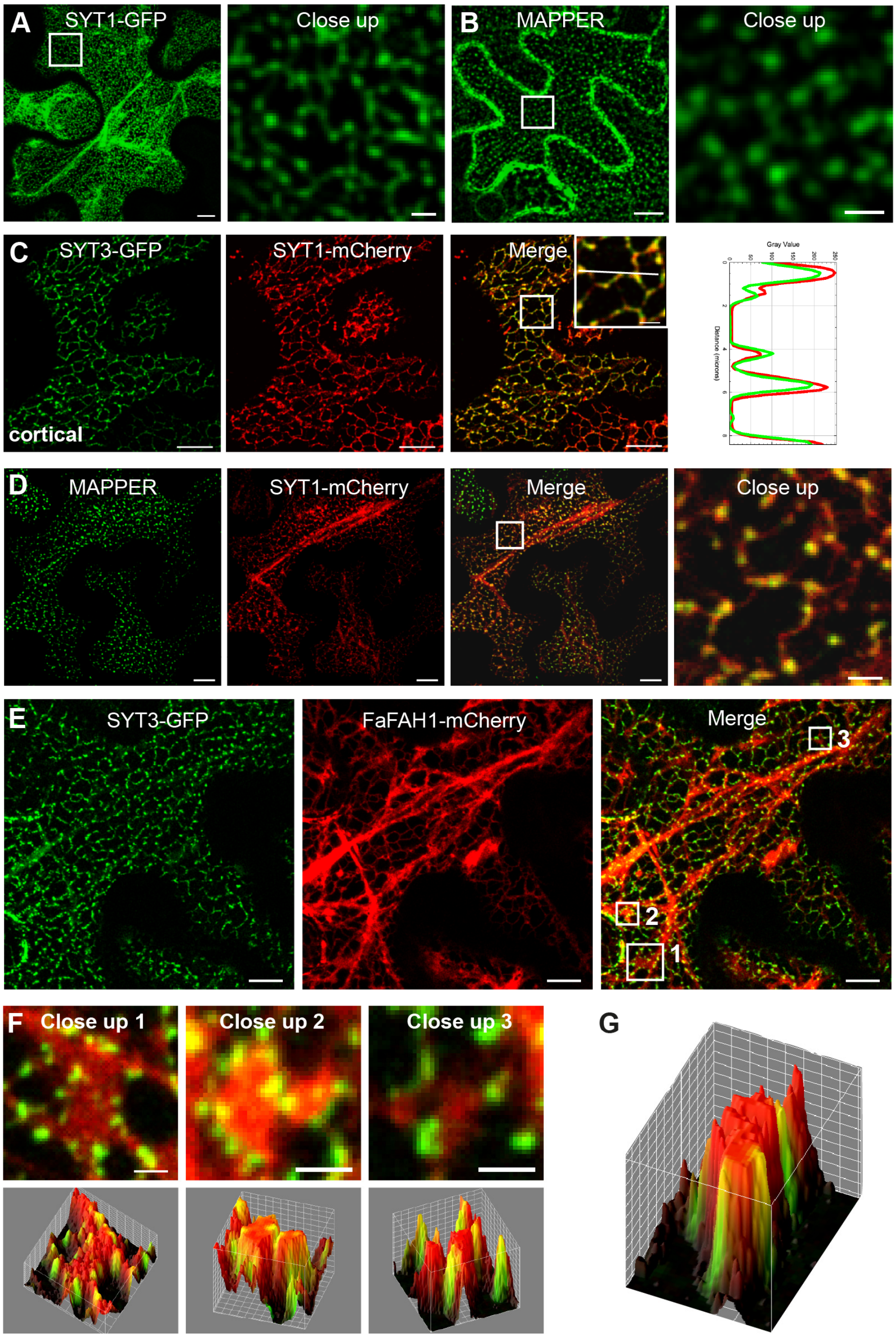


**Figure S4. SYT1 and SYT localize at ER-PM CS (related to Figure 3).**

**(A)** Confocal images of the cortical region of lower epidermal cells of *N. benthamiana* leaves transiently expressing *SYT1-GFP* 2. Boxed region is magnified in the close up.

**(B)** Confocal images of the cortical-to-equatorial region of lower epidermal cells of *N. benthamiana* leaves transiently expressing *MAPPER*. Boxed region is magnified in the close up.

**(C)** Confocal images of the cortical region of lower epidermal cells of *N. benthamiana* leaves transiently co-expressing *SYT3-GFP* with *SYT1-mCherry*. Images of the individual channels as well as the merged image are shown. The boxed region is magnified in the inset (close up). The intensity plot along the white line in close up view is shown.

**(D)** Confocal images of the cortical region of lower epidermal cells of *N. benthamiana* leaves transiently co-expressing *MAPPER* with *SYT1-mCherry*. Images of the individual channels as well as the merged image are shown. The boxed region is magnified in the close up.

**(E)** Confocal images of the cortical region of lower epidermal cells of *N. benthamiana* leaves transiently co-expressing *SYT3-GFP* with the ER marker *FaFAH1-mCherry*. Images of the individual channels as well as the merged image are shown. The boxed regions are magnified in Figure S4F.

**(F)** Close up views of the boxed regions in Figure S4E (upper panels) and 3D-surface intensity plots of the green and red channels (lower panel). Three D-surface intensity plots were obtained with the FIJI plugin *3D surface plot*. Min and max intensity values were adjusted in the interactive 3D surface plot viewer to improve visualization and to show that no green signal is observed in the interior or the red signals, only in the periphery.
**(G)** 3D-surface intensity plot of the green and red channels of the ER-sheet in the inset of Figure 3D. 3D-surface intensity plots were obtained with the FIJI plugin *3D surface plot*. Min and max intensity values were adjusted in the interactive 3D surface plot viewer to improve visualization and to show that no green signal is observed in the interior or the red signals, only in the periphery.
Scale bar 10 μm for all the wide-view images and 2 μm for all the close-up images.

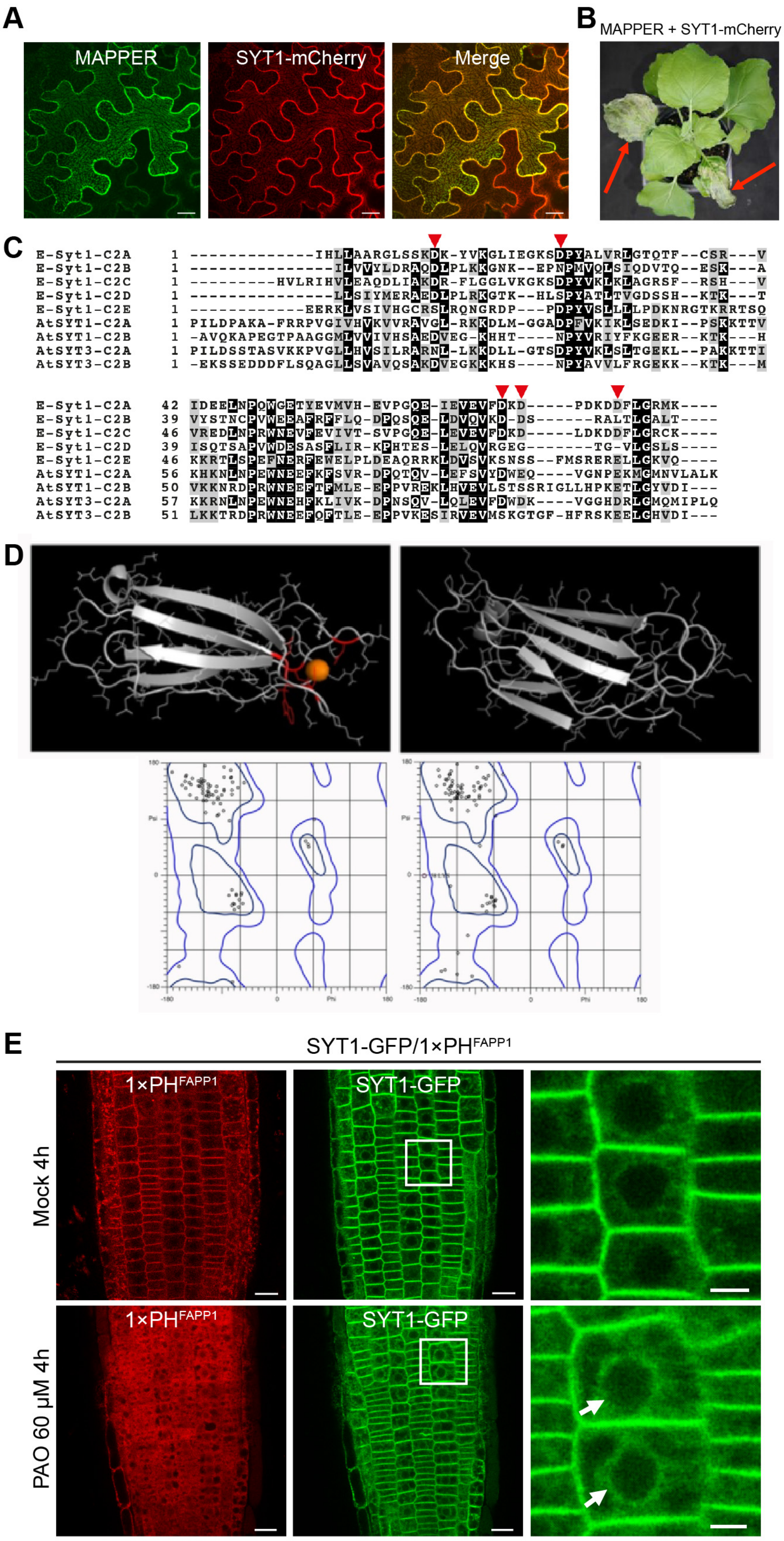


**Figure S5. SYT1 and SYT3 function as ER-PM CS tether and bind PI4P at the PM (related to Figure 4).**

**(A)** Confocal images of the cortical region of lower epidermis cells of *N. benthamiana* leaves transiently co-expressing *MAPPER* and *SYT1-mCherry*. In situations where both MAPPER and SYT1-mCherry were highly expressed, a massive expansion of ER-PM CS was observed. Scale bar 5 μm.
**(B)** A *N. benthamiana* plant in which two leaves have been co-infiltrated with *MAPPER* and *SYT1-mCherry*, rendering leaf necrosis (associated with high expression of both, MAPPER and SYT1-mCherry).

**(C)** Protein sequence alignment of C2 domains from Arabidopsis SYT1 and from *Homo sapiens* E-Syt1. The protein sequences were retrieved from the NCBI database. Multiple sequence alignment was performed using the Clustal W algorithm and formatted using the Box shade tool (http://www.ch.embnet.org/software/BOX_form.html). Black and gray boxes highlight identical and similar amino acids, respectively.
**(D)** Model of SYT3 C2 domains (upper panel) and analysis of torsional angles using Rampage (Lovell et al., 2003) and Ramachandran plots were generated (lower panel). As a result, very robust $C2A^{244\text{-}402}$ and $C2B^{403\text{-}540}$ 3D models with a high confidence match (>99%) were generated. Three $Ca^{+2}$ coordination pockets with a high confidence score (Cs >0.5) were identified for C2A2 while no $Ca^{+2}$ coordination pockets were identified for C2B.
**(E)** Images of roots after mock or PAO treatment. Plants are expressing 1×PH or SYT1-GFP. Scale bar, 10 µm and 2 µm for the close-up images

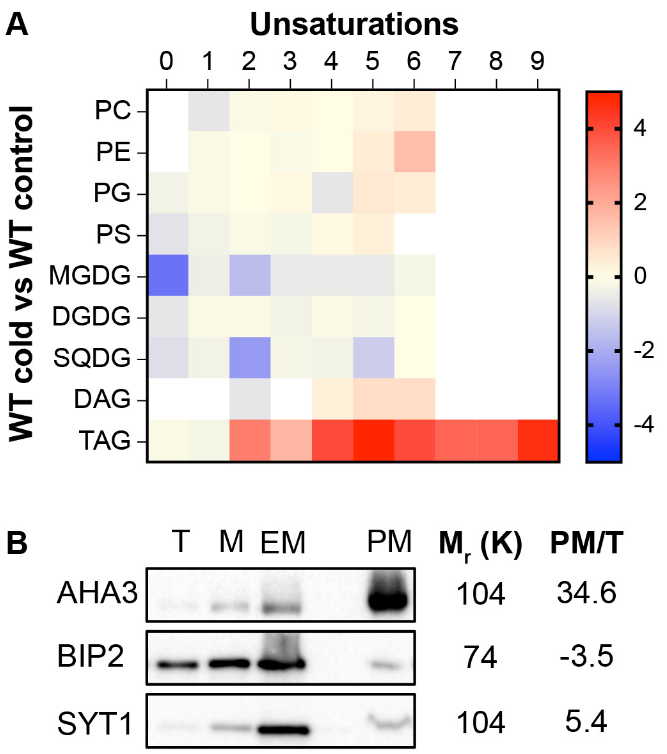


**Figure S6. Total lipid changes in WT plants after cold and analysis of PM fractions (related to Figure 6)**

**(A)** Global changes in the glycerolipids of Arabidopsis WT plants upon cold depending on the unsaturation grade of the acyl chain. The panel represents the fold change in the glycerolipidome of plants grown in control conditions against plants after a cold treatment. Scale code expresses fold changes in the relative peak area of the corresponding acyl lipids (n=3). PC, phosphatidylcholine; PE, phosphatidylethanolamine; PG, phosphatidylglycerol; PS, phosphatidylserine; LGPL, lysoglycerophospholipid; MGDG, monogalactosyldiacylglycerol; DGDG, digalactosyldiacylglycerol; SQDG, sulfoquinovosyldiacylglycerol; DAG, diacylglycerol; TAG, triacylglycerol.

**(B)** Quantification of protein abundance after the PM purification (see Methods). Total (T) crude homogenized tissue samples before purification, microsomes (M), final endomembranes (EM) and plasma membrane (PM) samples were processed by SDS–PAGE and immunoblotting with the indicated antibodies. Note the enrichment of PM protein (AHA3) and depletion of ER protein (Bip) in the PM fractions.

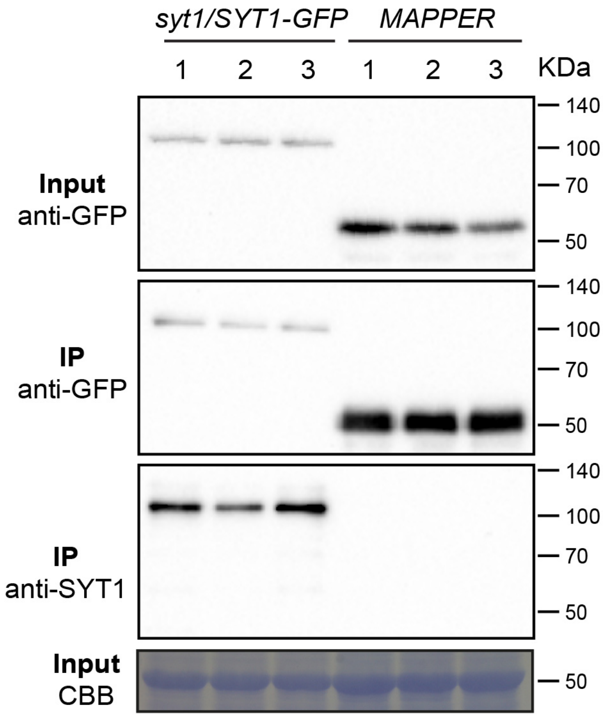


**Figure S7. SYT1-GFP and MAPPER do not associate at ER-PM CS (related to Figure 7),**
SYT1-GFP and GFP-MAPPER proteins were immunoprecipitated using anti–GFP-Trap beads. The non-denaturing zwitterionic detergent CHAPS was used for solubilizing the membrane proteins and breaking protein-protein interactions. Total (input) and immunoprecipitated (IP) proteins were separated by SDS-PAGE. GFP-tag proteins were detected with anti-GFP and SYT1 was detected with anti-SYT1 antibody, respectively. SYT1 protein did not co-immunoprecipitated with MAPPER. Equal loading was confirmed by Coomassie blue staining (CBB) of input samples. Molecular weight (kDa) marker bands are indicated for reference. Three biological replicates were analyzed.
